# Phylogenetic and metabolic diversity have contrasting effects on the ecological functioning of bacterial communities

**DOI:** 10.1101/839696

**Authors:** Constantinos Xenophontos, Martin Taubert, W. Stanley Harpole, Kirsten Küsel

**Affiliations:** Institute of Biodiversity, Friedrich Schiller University Jena, Jena 07743, Germany; German Centre for Integrative Biodiversity Research (iDiv) Halle-Jena-Leipzig, 04103 Leipzig, Germany; Department of Physiological Diversity, Helmholtz-Center for Environmental Research (UFZ), 04318 Leipzig, Germany; Institute of Biology, Martin Luther University of Halle-Wittenberg, 06120 Halle, Germany

**Keywords:** ecosystem functions, phylogenetic diversity, metabolic diversity, biodiversity, bacteria, synthetic communities, exoenzymes, community interactions

## Abstract

Quantifying the relative contributions of microbial species to ecosystem functioning is challenging, because of the distinct mechanisms associated with microbial phylogenetic and metabolic diversity. We constructed bacterial communities with different diversity traits and employed exoenzyme activities (EEAs) and total available carbon (TAC) from substrates as proxies of bacterial functioning to test the independent effects of these two aspects of biodiversity. We expected that metabolic diversity, but not phylogenetic diversity would be associated with greater ecological function. Phylogenetically relatedness should intensify species interactions and coexistence, therefore amplifying the influence of metabolic diversity. We examined the effects of each diversity treatment using linear models, while controlling for the other, and found that phylogenetic diversity strongly influenced community functioning, positively and negatively. Metabolic diversity, however, exhibited negative or non-significant relationships with community functioning. When controlling for different substrates, EEAs increased along with phylogenetic diversity but decreased with metabolic diversity. The strength of diversity effects was related to substrate chemistry and the molecular mechanisms associated with each substrate’s degradation. EEAs of phylogenetically similar groups were strongly affected by within-genus interactions. These results highlight the unique flexibility of microbial metabolic functions that must be considered in further ecological theory development.

## Introduction

Bacterial activities are driving ecosystem functioning (Finlay, Maberly and Cooper 1997) and recent research has highlighted the contribution of microbial diversity to the biological processes occurring in ecosystems (Petchey and Gaston 2006; Cardinale *et al*. 2012; Galand *et al*. 2018; Le Bagousse-Pinguet *et al*. 2019; Johnson and Pomati 2020). As such, a direct assessment of metabolic diversity is a great predictor of community ecosystem functions (Mokany, Ash and Roxburgh 2008; Salles *et al*. 2009; Krause *et al*. 2014). Microbial metabolic traits can be determined based on, e.g., functional gene-targeting microarrays and metabolic activity assays (Litchman and Klausmeier 2008; Naeem *et al*. 2009; Paula *et al*. 2014; Wang *et al*. 2017). Such assays have shown that metabolic diversity is positively correlated with ecologically important community functions, such as organic matter decomposition and CO2 production (Rivett *et al*. 2016; Evans *et al*. 2017). Microbial functions like organic matter degradation are often conveyed by exoenzymes. The production and release of exoenzymes is an important mechanism that bacteria use to access exogenous, complex carbon and energy sources. Inevitably, exoenzymes are shared among different species (Drescher *et al*. 2014; Smith and Schuster 2019), and thus their production is expected to be affected by community species interactions (Folse and Allison 2012). Exoenzyme activities (EEAs) can also link multiple bacterial metabolic processes to ecosystem functions like carbon cycling (Malik *et al*. 2019). Moreover, metabolic traits affect fitness, as they relate to energy acquisition and expenditure, placing them under the influence of strong evolutionary selection processes. If we aim to improve our understanding of how metabolic diversity influences microbial community ecosystem functioning, the evolutionary history and relationship of microbes must be considered.

Phylogenetic relationships are usually assessed using neutral genetic markers like the bacterial 16S rRNA gene. But as a neutral marker it is not subject to selective pressures. So ecologists have been debating the usefulness of phylogenetic diversity in the study of ecosystem functioning, as studies have yielded mixed results (Cadotte *et al*. 2009; Flynn *et al*. 2011; Alexandrou *et al*. 2015; Gerhold *et al*. 2015; Venail *et al*. 2015). Nevertheless, closely related bacterial species are often more likely to have similar metabolic traits (Martiny, Treseder and Pusch 2013; Goberna and Verdú 2016). Phylogenetic diversity is a good predictor for metabolic functions with a strong phylogenetic clustering: For example, ammonia oxidation is restricted to a group of chemolithoautotrophic bacteria (Kowalchuk and Stephen 2001), traced in deep phylogenetic branches. Yet, especially for broad, simple functions such as carbon assimilation from amino acids and simple sugars (Martiny, Treseder and Pusch 2013), it is a poor predictor of ecological functioning (Srivastava *et al*. 2012; Martiny, Treseder and Pusch 2013; Goberna and Verdú 2016). Nevertheless, under certain circumstances, phylogenetic relationships can predict growth rates (Morrissey *et al*. 2016), species interactions via ecological networks (Srivastava *et al*. 2012) and species coexistence (Germain, Weir and Gilbert 2016). Hence, we can link phylogenetic relatedness to properties that are under natural selection and therefore contribute to ecological functioning (Morrissey *et al*. 2016).

The precise connection behind the phylogenetic and metabolic diversities is still unclear, because the underlying mechanisms are either unknown or very complex (Losos 2008). Since phylogenetic diversity is connected with species coexistence and interactions, which should interact with metabolic diversity (Pascual-García, Bonhoeffer and Bell 2020) and possibly modify or intensify the effects metabolic diversity has on community functioning (Nealson and Popa 2005). For the further development of micro-ecological theory (Krause *et al*. 2014; Shade *et al*. 2018) and biodiversity research (Flynn *et al*. 2011; Roger *et al*. 2016), potential interactions between phylogenetic and metabolic diversity have to be identified to understand their influence on community ecosystem functions (Thompson, Davies and Gonzalez 2015).

Here, we aim to disentangle this interaction by using synthetic microbial communities in a multifactorial experimental design, where we control for each diversity aspect while studying the other’s effect on community ecological functions. We used groundwater bacterial isolates to form simplified four-member communities with different phylogenetic and metabolic diversity treatments, allowing us to test how these two aspects of biodiversity interact as they shape community ecological functions. EEA assays were used for the metabolic classification of bacterial isolates and to assess the functioning of synthetic communities. We hypothesised that 1) when controlling for phylogenetic diversity, metabolic diversity has a positive effect on community ecological functions. In contrast, 2) when controlling for metabolic diversity, phylogenetic diversity, based on a neutral marker, should have no effect on community ecological functions. Finally, 3) the effect of metabolic diversity is proportionally stronger in communities that are phylogenetically similar, as phylogenetically dissimilar communities hold species with less niche overlap and therefore negative interactions like competition are weaker.

## Material and Methods

### Selection of bacterial isolates and culture setups

The twenty-seven bacterial strains used in this study (Appendix S1: Table S1) were isolated from pristine groundwater (Geesink *et al*. 2018), obtained from the Hainich Critical Zone Exploratory located in Thuringia, Germany (Küsel *et al*. 2016). The isolates were selected from a pool of more than 340 groundwater isolates as representatives of the most abundant taxonomic families within the total groundwater bacterial population, based on 16S rRNA gene MiSeq amplicon sequencing data (Kumar *et al*. 2017; Schwab *et al*. 2017), throughout the taxonomic classes of *Actinobacteria, Bacilli, Flavobacteriia, Sphingobacteria, Alpha*- and *Gammaproteobacteria*.

The isolates were maintained on S2P medium agar (Geesink *et al*. 2018). Before each of the experimental assays, isolates were cultured aerobically in 20 ml of liquid S2P medium, in cap-perforated 50 ml Falcon tubes, horizontally shaken (125 rpm) at room temperature, for 48 h to early stationary phase. Growth in the liquid cultures was being determined via optical density measurements at a wavelength of 600 nm (Synergy™ H4 Hybrid Multi-Mode Microplate Reader, BioTek, Germany).

### Molecular analysis and taxonomic classification of bacterial isolates

DNA from bacterial isolates was extracted with GenElute™ Bacterial Genomic DNA Kit (NA2110-1KT, Sigma-Aldrich, Germany). Sterile inoculation loops were used to gather 1 μl-worth of biomass from the agar medium cultures, for DNA extraction. For the 16S rRNA gene amplification, the primers 8F (5’-AGA GTT TGA TCC TGG CTC AG-3’) (Turner *et al*. 1999) and 1492R (5’-GGT TAC CTT GTT ACG ACT T-3’) (Turner *et al*. 1999) were used. PCR was performed with a PEQSTAR thermocycler (PEQLAB, Germany) using the following cycling conditions: initial denaturation at 95 °C for 10 min, 30 cycles of 95 °C (denaturation), 55 °C (primer annealing) and 72 °C (elongation) for 30 sec, 45 sec and 60 sec, respectively, followed by 10 min at 72 °C (final elongation). DNA purification and sequencing of the 1484bp-length PCR products was carried out by Macrogen (Amsterdam, Netherlands) with the 27F (5’-AGA GTT TGA TCM TGG CTC AG-3’) primer.

The sequences were assigned to their closest reference relative using BLASTn (Zhang *et al*. 2000) (v2.8.0+) and cross-referenced with SILVA Incremental Aligner (SINA) (Pruesse, Peplies and Glöckner 2012) (v1.2.11). A rooted maximum-likelihood phylogenetic tree for the 16S rRNA gene taxonomic classification of the organisms was constructed with MEGA X (Kumar *et al*. 2018) (v10.1.5).

### Exoenzyme activity (EEA) assays

EEA assays for each bacterial isolate were performed using eleven substrates (Appendix S1: Table S2) (Sigma-Aldrich, Germany), each carrying a fluorescent moiety allowing spectrophotometric measurements of activity using a microplate reader (Synergy™ H4 Hybrid Multi-Mode Microplate Reader). The sensor was set to detect fluorescence at wavelengths 360/9 nm for excitation and 450/9 nm for emission.

Prior to the EEA assays, the optical density of the bacterial cultures was determined spectrophotometrically (600 nm). For the EEA assays, a modified version of the protocols from Frossard *et al*. (Frossard *et al*. 2012) and Rivett *et al*. (Rivett *et al*. 2016) was used here, as follows: a black 96-well plate setup, with 100 μl HEPES buffer at 100 mM stock concentration in each well, was inoculated with 100 μl of bacterial culture and to each test well 50 μl of substrate solution at 40 μM final concentration was added. Negative controls consisted of wells containing 100 μl of sterile S2P liquid medium and wells with just 100 μl sterile Milli-Q water (Merck Millipore, Germany). Fluorescence standards for the MUB substrates were measured using 4-Methylumbelliferone with final concentrations of 10 to 0.3125 μM by halving dilutions. Fluorescence standards for the AMC substrates were measured using 7-Amino-4-methylcoumarin with final concentrations of 2 to 0.0625 μM by halving dilutions. The standards were used to transform relative fluorescence units (RFU) to concentration of degraded substrate. Each assay plate setup held technical triplicates for the cultures, controls and standards. Triplicates were averaged and their standard deviation propagated for the data analyses. Background fluorescence, measured in negative controls with either substrate solution or Milli-Q water, was subtracted from the respective assay plates. Measurement of the assay plates begun immediately after the substrate addition and continued for 90 min, at 2.5 min measurement intervals.

### Determining phylogenetic and metabolic diversity of isolates

To examine effects of diversity on the functioning of synthetic bacterial communities, the 16S rRNA gene taxonomy and EEAs of the twenty-seven bacterial isolates were used as the basis of the community traits: phylogenetic and metabolic diversity. Using seventeen of the tested isolates, we formed four trait-groups, each consisting of five isolates, being either phylogenetically and metabolically similar (Psim-Msim), phylogenetically similar but metabolically dissimilar (Psim-Mdis), phylogenetically dissimilar and metabolically similar (Pdis-Msim) as well as phylogenetically and metabolically similar (Pdis-Mdis). In total, the isolates covered ten taxonomic families of five classes (Appendix S1: Figure S1). In the Psim groups, isolates were affiliated with the same genus; Psim-Msim group held *Flavobacteria* isolates and Psim-Mdis held *Pseudomonas* isolates. Pdis groups were formed so that all members of a group were related to different families. Subsequently, Psim and Pdis group phylogenetic diversity quantitative measures were obtained using the “*Compare within group mean distance*” function of MEGA X (Kumar *et al*. 2018) (v10.1.5) and verified to be statistically insignificant for Psim communities and significantly different for Pdis communities using paired t-tests (Appendix S1: Table S3). Hierarchical clustering based on Euclidean distance was used to quantise functioning of the total EEAs for each isolate (Figure 1) and categorise them as metabolically similar or dissimilar. The clustering analysis was performed in R (v. 3.6.1) (R Core Team 2019) using the core stats functions *dist(x, method = “euclidean”, p = 2)* to calculate the Euclidean dissimilarity distance matrix (Borg and Groenen 2005), followed by *hclust(d, method = “ward.D”)* to hierarchically rank and cluster the EEA dissimilarities between the bacterial isolates using the Ward hierarchical clustering method (Ward 1963; Murtagh and Legendre 2014). Msim groups were formed from isolates of the same cluster, Mdis groups were formed so that each member of the group belonged to a different metabolic cluster (Figure 1). Quantitative measures of principal component analysis (PCA) based on the metabolic traits of the isolates confirmed the higher metabolic diversity between isolates of Psim-Mdis and Pdis-Mdis compared to group Psim-Msim and Pdis-Msim (Appendix S1: Figure S2).

**Figure 1.**
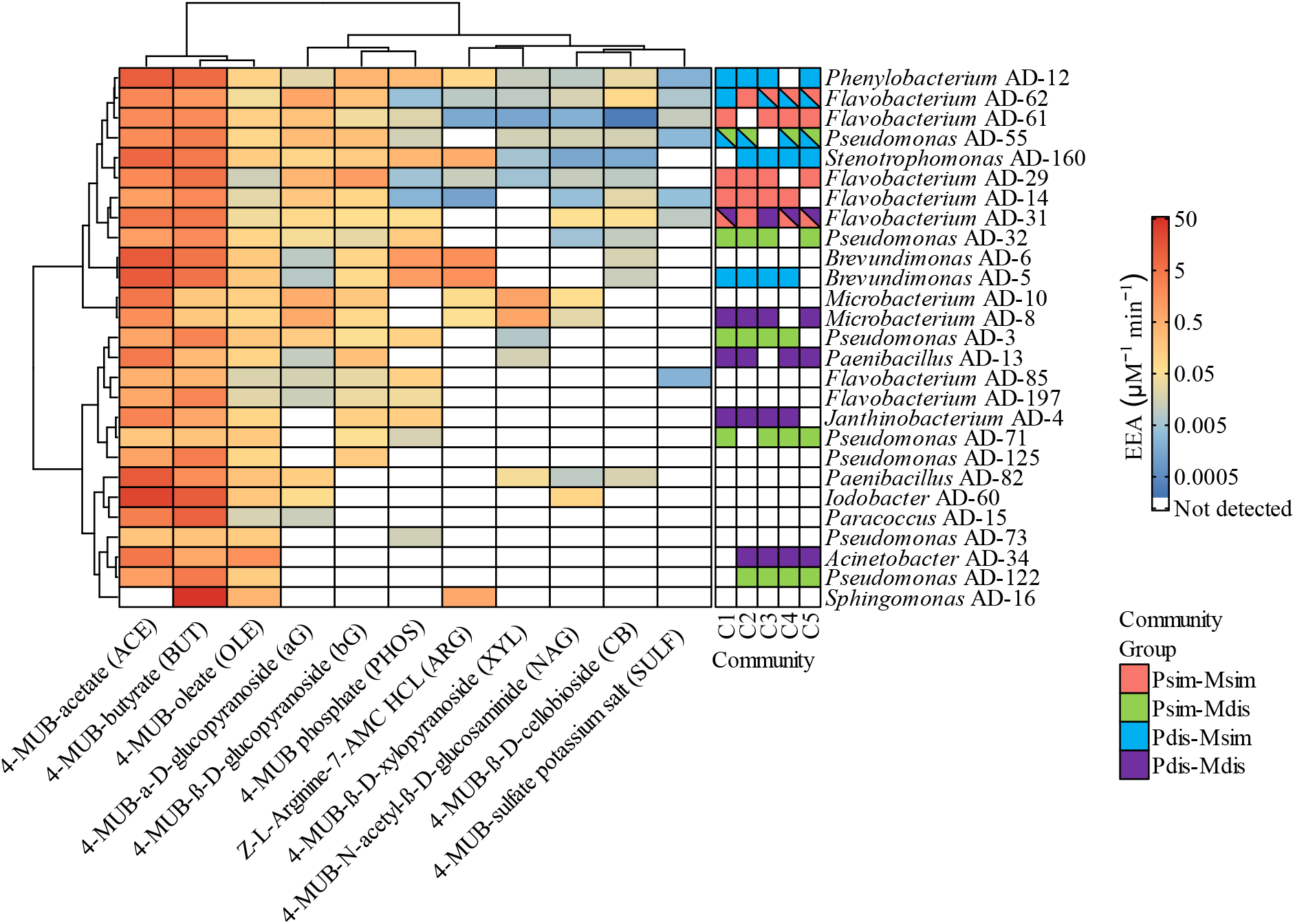
Metabolic and phylogenetic classification and grouping of groundwater isolates. EEAs are shown for twenty-seven groundwater isolates incubated with eleven fluorescent substrate analogues. The heatmap scale shows the concentration of substrate that was broken down in the EEA assays. Empty (white) boxes denote activity outside the detectable range. The left dendrogram shows clusters of isolates with similar metabolic traits based on Euclidean hierarchical diversity distances. The topmost dendrogram shows hierarchical clustering of substrates that produced similar activity patterns in the assays. The assignment of isolates to the communities of each treatment group is depicted to the right of the heatmap. Doublecoloured boxes indicate isolates that were assigned to more than one group. In the legend, the colours indicate the community group and the phylogenetic (P) and metabolic (M) traits of the communities are indicated as similar (sim) or dissimilar (dis).

### Bacterial community setups and experimental design

For each of the four trait groups with five isolates (Psim-Msim, Psim-Mdis, Pdis-Msim, Pdis-Mdis), we designed five distinct synthetic communities of bacterial species by omitting one isolate each per community. This resulted in twenty communities, five for each trait group, identified as C1 to C5. We confirmed the phylogenetic diversity between the communities via the quantitative estimates of isolate phylogenetic distance and their metabolic diversity based on the Euclidean distance matrix of isolate EEAs as previously mentioned (Appendix S1: Table S3). The phylogenetic and metabolic diversity values for each community showed a distinct differentiation of the communities from each of the four groups, confirming that we successfully separated the bacterial isolates and communities into the four intended groups (Figure 2).

**Figure 2.**
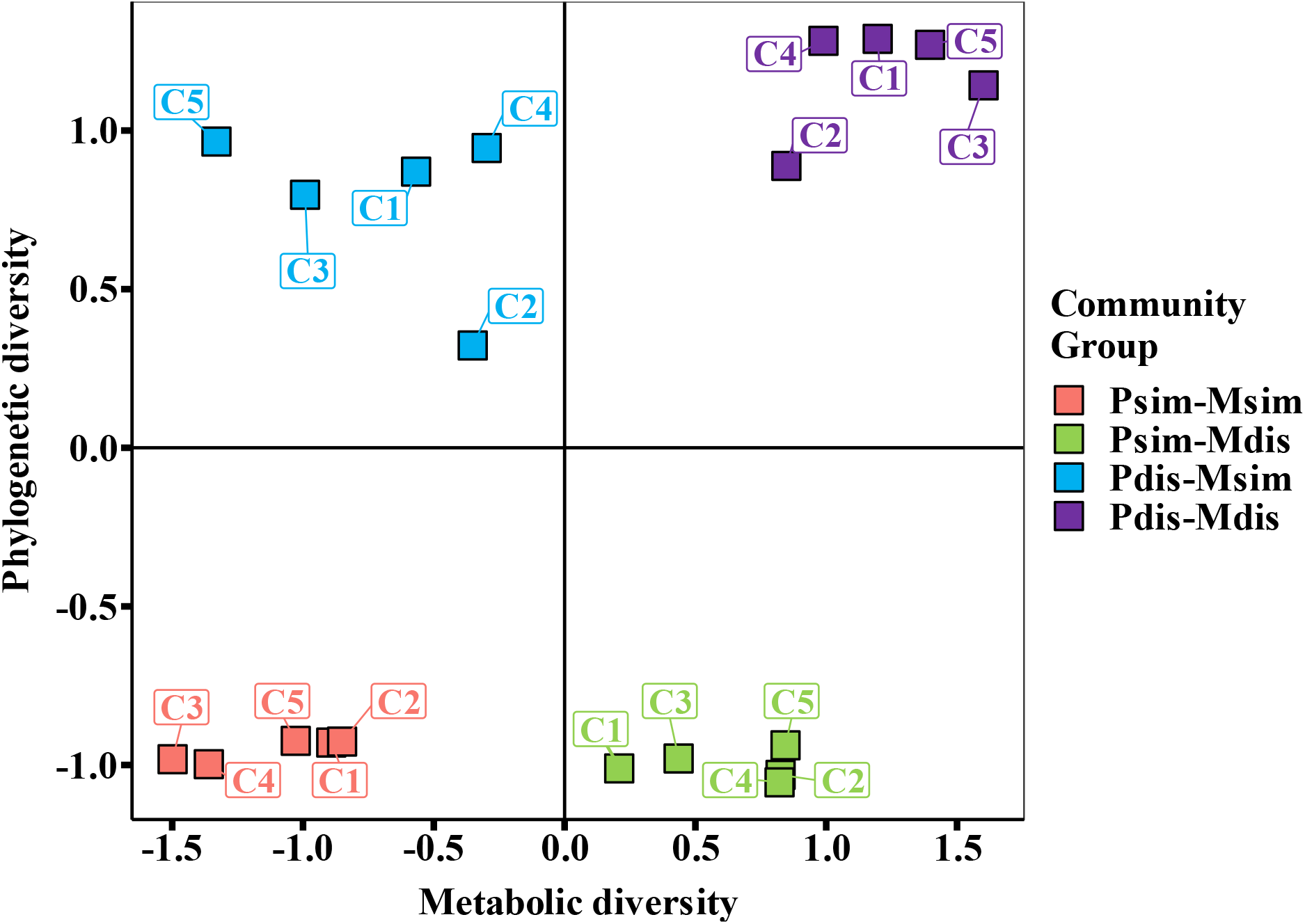
Individual communities plotted against the phylogenetic and metabolic diversity within and between groups. Colours indicate the community groups; red for Psim-Msim group, green for Psim-Mdis group, blue for Pdis-Msim group and purple for Pdis-Mdis group. Phylogenetic and metabolic diversity metrics have been z-score normalised. Variation of diversity scores exists within groups, but the groups are cleanly separated from each other (see Appendix 1: Table S3 for statistical significance).

We grew cultures of each isolate in liquid S2P medium as described above. Optical density (600 nm) was used to calculate the required final volume for each isolate, so that all members of each community were to be introduced into the communities at the same amount of biomass. The communities were established in cultures by mixing a predetermined amount of liquid isolate culture for a total volume of 2.5 ml. The community cultures were incubated in deep 96-well plates (2.5 ml volume), at room temperature and sealed with rayon film seals (391-1262, VWR, Germany) to allow gas exchange. The EEAs of the communities were assessed after 0 h, 6 h and 24 h of incubation. Additionally, at the 0 h and 24 h marks, a 200 μl culture sample was pelleted by centrifugation (17.000 × g, 5 min) and stored at −80 °C, for subsequent molecular analysis.

### Ecological functioning data analysis

As a measure of multifunctionality, we used the total available carbon (TAC) from each substrate for a direct comparison of communities. This was done by multiplying the total number of carbon atoms in each substrate (excluding the fluorescent moiety) with the molar amount of degraded substrate. TAC was summed as a unified measure of ecological functioning for each isolate and community. 4-MUB phosphate (PHOS) and 4-MUB sulfate (SULF) were excluded because they contained no carbon atoms.

In addition, the effect of the diversity treatments on each individual EEA was determined. The communities constructed in this study were composed out of different isolates with widely differing metabolic activities. A comparison of EEAs between communities alone would thus only reflect the differences between the constituent isolates. Therefore, EEA differences of the communities were normalized to the respective isolates and expressed as proportional changes of EEA. This was done with the following steps (visualised in Appendix S1: Figure S3): The measured EEAs of the isolates within each community group were used to calculate expected EEAs for each of the group’s communities. The expected EEAs were compared to the measured EEAs of the experimental communities and the result was represented as % proportional changes of EEAs. This resulted in values from −100% to 385%.

To evaluate the significance of phylogenetic diversity, metabolic diversity was controlled by subsetting into metabolically similar and dissimilar communities. Likewise, to examine the significance of metabolic diversity, phylogenetic diversity was controlled for in the same way. These analyses were performed for the 0 h timepoint data, using linear regression models in R. For the linear models, the TAC response variable data, were log2-transformed. To also enable logarithmic scaling of the EEA % proportional change data, 101% was added to all datapoints, to avoid negative values and zeroes, since the smallest values were −100%; As the TAC values were all > 0, this was not necessary. To avoid pseudoreplication in the models, the 11 EEA % proportional change datapoints of each community were averaged in the diversity treatments. For example, group Psim-Fsim community C1 had eleven proportional change values, one for each substrate, that were averaged into one. This was repeated for all communities in all group and resulted in a total of 20 datapoints. The group Pdis-Mdis showed no activity in the SULF assay, and so this assay was excluded from the models. For the TAC data, 20 datapoints were also used, one calculated value per community. Through subsetting the predictor variable data, to control for each diversity effect, 10 datapoints were used for each model, for both EEA % proportional change data as well as the TAC data. Furthermore, predictor variables, phylogenetic and metabolic diversity metrics, were z-score normalised, for the linear regression models, to make them comparable.

For the analyses for the influences of substrate biochemistry on the diversity treatment effects, the EEA% proportional change data were not subset by diversity treatment but by each substrate. Therefore, the EEA % proportional change response variable for each substrate model was 20 datapoints or 15 for SULF. Following that, ANOVA of linear regression models was performed as described above.

### Effects of diversity in EEA changes over 24 h

Time is a crucial driver of bacterial interactions and therefore we evaluated how the EEA proportional changes of communities developed over 24 h, by comparing the 0 h timepoint to the 6 h and 24 h timepoint data. This was done for all 20 communities of all 4 diversity treatment groups, and for each substrate, the average of each group was determined. The significance of temporal changes was tested using the Student’s t-test (2-tailed) by comparing the EEA proportional change of all communities in a treatment group at 0 h to all EEA proportional changes of the same group for timepoints 6 h and 24 h, for each individual substrate. False discovery rate was controlled with the Benjamini–Hochberg procedure (McDonald 2014).

### Substrate acclimation in EEA assays

To study influences of acclimation of isolates and communities to the added substrates on the observed effects of diversity treatments on community ecological functions, we added a pre-incubation period to the EEA assays. This was performed only with the Psim-Mdis group. After mixing of the communities, substrates were added after 0 h and 6 h to isolates and communities, and EEA assays were performed as described above (no-acclimation treatments). In addition, isolates and communities with substrates added 0 h after mixing were incubated for a total of 4.5 h, before another series of EEA assays were conducted (acclimation treatments).

Assay duration was 90 min, as in previous assays. To achieve a direct comparability of the activities measured with and without acclimation, the determined EEAs were normalized to optical density of isolates and communities, which was measured before each assay. Following that, the change in EEAs from 0 h to 4.5 h and 6 h was compared between isolates and communities.

### 16S rRNA gene amplicon sequencing of synthetic communities

Community DNA was extracted as described above. DNA concentration was measured spectrophotometrically (NanoDrop 1000, PEQLAB, Germany) and samples were stored at −20 °C.

For Illumina MiSeq amplicon sequencing of 16S rRNA genes, PCR was performed using the primer set Bakt_341F (5’-CCT ACG GGN GGC WGC AG-3’) (Herlemann *et al*. 2011) and Bakt_805R (5’-GAC TAC HVG GGT ATC TAA TCC-3’) (Herlemann *et al*. 2011) targeting the V3-V4 hypervariable region of the rRNA gene. DNA amplification was carried out with a PEQSTAR 2X thermocycler (PEQLAB, Germany) in the following conditions: initial denaturation at 95 °C for 15 min, 30 cycles of 94 °C (denaturation), 55 °C (primer annealing) and 72 °C (elongation) at 45 sec for each step, followed by 10 min at 72 °C (final elongation). The PCR DNA products were purified using NucleoSpin Gel and PCR Clean-up (Ref. 740609, Macherey-Nagel, Germany) following the manufacturer’s protocol. The final step, DNA recovery from the columns, was performed twice for a total volume of 60 μl purified PCR product samples.

MiSeq-generated sequence data were analysed using mothur (Schloss *et al*. 2009) (v.1.39.1; mothur.org) along with the mothur MiSeq SOP developed by Kozich *et al*. (Kozich *et al*. 2013) (accessed November 19th 2018). Paired-end reads were combined and sequences below 400 bp or above 429 bp as well as sequences with ambiguous bases or homopolymer runs above 8 bp were removed. Sequences were aligned to the SILVA reference database release SSU 132 (Quast *et al*. 2012) and clustered when differed in four or fewer bases. Chimeric sequences were removed with uchime (Edgar 2016) using the GOLD database (Mukherjee *et al*. 2019) as a reference. OTUs were binned from sequences with a 3% identity cut-off and were classified using the same SILVA database. Raw data have been deposited in the European Nucleotide Archive under the study with accession number PRJEB34832 (samples ERR3588683 - ERR3588722). The amplicon sequencing data were used to examine shifts in the Pdis community species relative abundances that might explain changes in the ecological functioning of communities that cannot be attributed to the diversity treatments.

## Results

### Effects of diversity on community ecological functions

We examined the absolute level of functioning of communities, using TAC turned over by EEA in each community as a unified measure (Figure 3; Appendix S1: Table S4). Linear regression analyses revealed that when controlling for metabolic diversity, phylogenetic diversity had a significant negative effect on TAC in metabolically similar communities (−0.305 ± 0.102, estimate ± std. error, Adj. R^2^ = 0.467, p = 0.018) (Figure 3A). This shows that communities with phylogenetically diverse isolates were able to acquire less carbon than communities with phylogenetically similar isolates, when these isolates had a high metabolic similarity. For metabolically dissimilar communities, phylogenetic diversity had a positive effect (0.313 ± 0.105, estimate ± std. error, Adj. R^2^ = 0.467, p = 0.018). While controlling for phylogenetic diversity, metabolic diversity had a negative effect in phylogenetically similar communities (−0.611 ± 0.151, estimate ± std. error, Adj. R^2^ = 0.632, p = 0.004) (Figure 3B), but no effect was observed for phylogenetically dissimilar communities (0.002 ± 0.115, estimate ± std. error, Adj. R^2^ = −0.125, p = 0.986).

**Figure 3.**
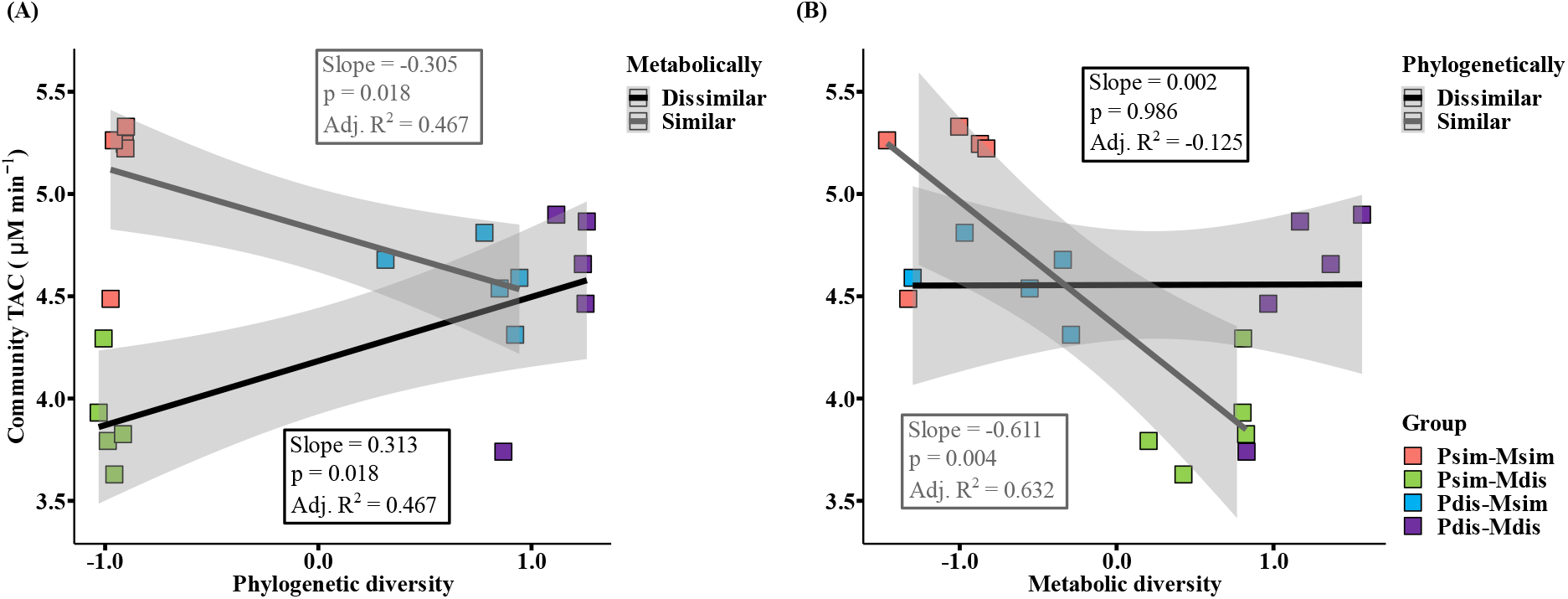
Effects of phylogenetic diversity (A) and metabolic diversity (B) on the total available carbon (TAC) of communities. Black regression lines show the effect of each treatment when controlling for the dissimilar groups of the second treatment. Grey regression lines show the effect of each treatment when controlling for the similar groups of the second treatment. Statistical significance for each linear regression is shown in each plot, with the respective coloured text (grey or black). Coloured squares indicate community groups: Psim-Msim in red, Psim-Mdis in green, Pdis-Msim in blue and Pdis-Mdis in purple. The y axes have been log2 transformed and the x axes have been z-scale transformed.

In addition to TAC, similar trends were also observed when performing linear regression analyses on community EEA changes (Appendix S1: Figure S4, Table S5). For metabolically similar communities, phylogenetic diversity had a significant negative effect on the EEAs (−0.279 ± 0.121, estimate ± std. error; Adj. R^2^ = 0.325; p = 0.050) when controlling for metabolic diversity (Appendix S1: Figure S4A). In contrast, for metabolically dissimilar communities, a significant positive effect of phylogenetic diversity was observed (1.094 ± 0.231, estimate ± std. error, Adj. R^2^ = 0.704, p = 0.002). While controlling for phylogenetic diversity phylogenetically similar communities exhibited a strong negative relationship, (−1.535 ± 0.366, estimate ± std. error, Adj. R^2^ = 0.648, p = 0.003), while phylogenetically dissimilar communities’ relationship was below the significance threshold (−0.121 ± 0.113, estimate ± std. error, Adj. R^2^ = 0.017, p = 0.314) (Appendix S1: Figure S4B). Hence, biodiversity effects in the carbon acquisition potential of these communities were also reflected on the community EEAs.

Interestingly, we furthermore observed that EEA changes of the synthetic four-species communities exhibited strong differences between substrates (Figure 4; Appendix S1: Figure S5). EEA changes showed sharp decreases, down to −100%, and increases, up to 385%, at 0 h after mixing. Substrates with glycosidic bonds like 4-MUB-N-acetyl-β-D-glucosaminide (NAG), 4-MUB-β-D-cellobioside (CB) and 4-MUB-β-D-xylopyranoside (XYL) exhibited the highest relative increases in EEA (see Appendix S1: Table S2 for complete list of substrate abbreviations). The EEA changes for each one substrate within a treatment group, however, were typically consistent (< 50% EEA proportional change from the average) for 33 of the 44 (75%) community-substrate combinations. This indicated that, in most cases, community behaviour was not strongly influenced by the excluded fifth isolate of each group. The consistency was more prominent for substrates with ester bonds like 4-MUB acetate (ACE), 4-MUB-butyrate (BUT) and 4-MUB oleate (OLE). A complete overview of cleaved substrate concentrations for isolates, communities and estimated community EEAs for each treatment group, can be found in the supplement (Appendix 1: Tables S6, S7, S8, S9). When comparing EEA changes between the groups (Appendix S1: Figure S5B-E), we observed that Pdis communities had relatively small to no changes in EEA: 76% of community-substrate combinations of group Pdis-Msim and Pdis-Mdis remained between +50% and −50% EEA proportional change. Contrarily, communities of Psim groups differed greatly in activities compared to their constituent isolates. Group Psim-Msim exhibited most of the largest EEA increases (up to 330%) with 27% of the group’s 55 community-substrate combinations measuring > +50% EEA proportional change. Group Psim-Mdis communities recorded the greatest amount of negative EEA proportional changes (53% of the group’s 55 community-substrate combinations having < −50% EEA proportional change) with the maximum EEA proportional change increase at only 5.4%. These results show that phylogenetically similar communities exhibited more variable and genus-specific responses than phylogenetically dissimilar communities.

**Figure 4.**
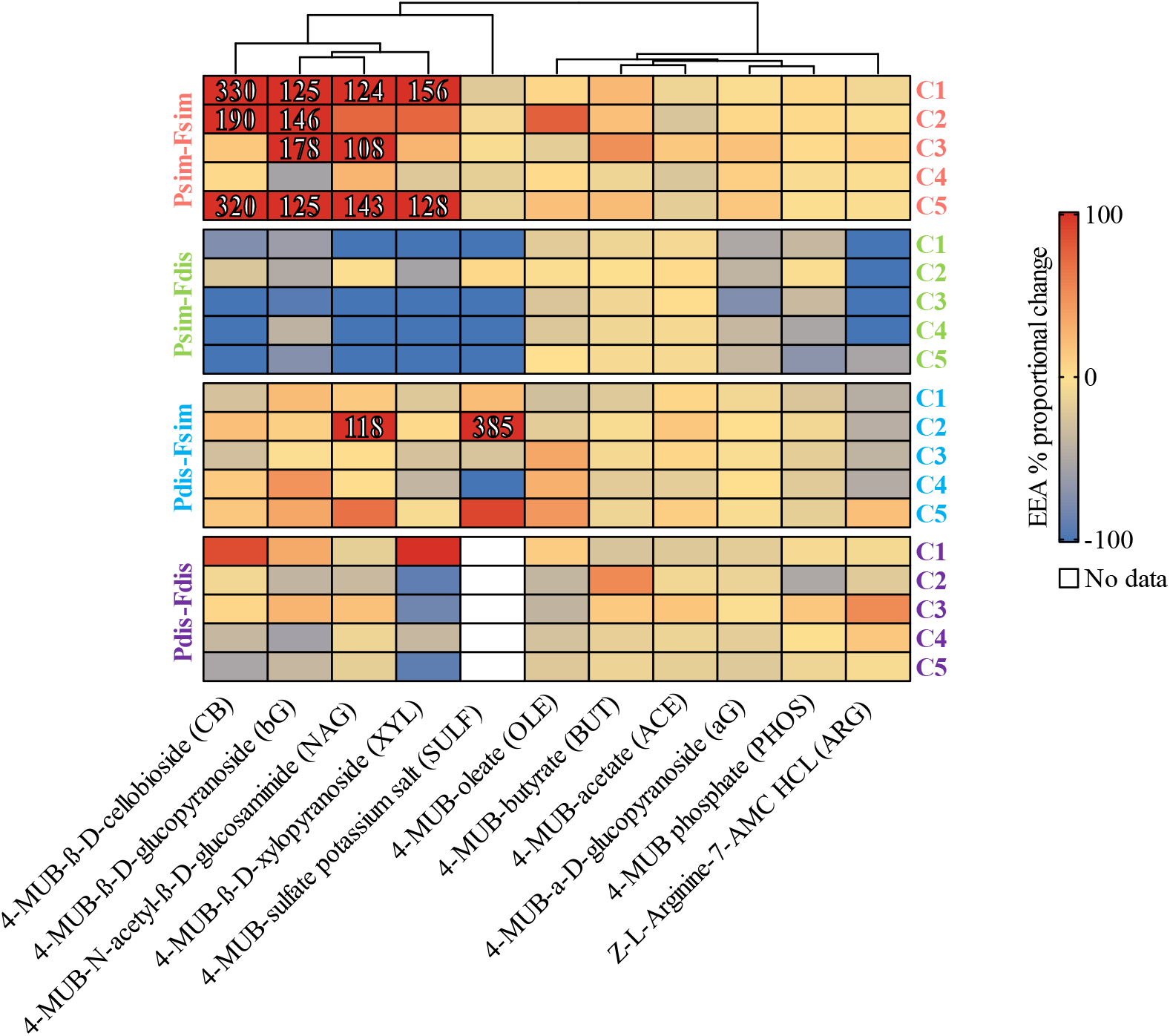
Heatmap of community EEA % proportional changes. The heatmap scale depicts proportional changes from −100 to 100%. Communities that exhibited changes over 100% have their percent change value shown in the heatmap matrix. Red colours depict a higher activity of the community; blue colours depict a lower activity of the community than estimated, based on the activities of the individual isolates. White boxes show measurements outside the detection limits. On the right, the label for each row indicates the community and on the left the group from where the activities are taken from; red for Psim-Msim group, green for Psim-Mdis group, blue for Pdis-Msim group and purple for Pdis-Mdis group communities. Columns indicate the substrates used in the EEA assays. The dendrogram depicts hierarchical clustering of each substrate’s activities, showing groups of substrates with similar activity responses.

### Chemical structure differences of substrates influence the strength of diversity effects

Investigating the datasets for all substrates individually using linear models, we observed broad differences in the relationships of EEAs to the metabolic and phylogenetic diversity between them (Appendix S1: Table S10). When controlling for substrates, the diversity treatments have a consistent pattern of influence, positive for phylogenetic diversity and negative for metabolic diversity (Figure 5). For example, CB showed a significant positive relationship with phylogenetic diversity (1.029 ± 0.444, estimate ± std. error; Adj. R^2^ = 0.520, p = 0.034) and for metabolic diversity, a significantly negative relationship (−1.604 ± 0.441, estimate ± std. error; Adj. R^2^ = 0.520, p = 0.002). The other two polysaccharide-analogue substrates, XYL and NAG, behaved similarly to CB (Appendix S1: Table S10). We observed that the strength of the effect was related with the chemical structure of the substrates (Figure 5). Esters (ACE, BUT, OLE) exhibited no significant relationship with diversity treatments, with linear slopes ranging from −0.095 to 0.038 for phylogenetic diversity and −0.227 to −0.007 for metabolic diversity. Monosaccharide (aG, bG) linear slopes ranged from 0.211 to 0.368 and −0.760 to −0.336, a relationship that was overall similar but weaker than for the polysaccharides (CB, XYL, NAG), whose linear slopes were 1.029 to 1.470 and −2.019 to −1.604 for phylogenetic and metabolic diversity, respectively. On average, by controlling for substrates alone, we observed that phylogenetic diversity had a positive effect on EEA changes (0.643 ± 0.193, estimate ± std. error; Adj. R^2^ = 0.645, p = 0.004) (Table S10). Contrastingly, metabolic diversity had a negative effect (−0.897 ± 0.191, estimate ± std. error; Adj. R^2^ = 0.645, p = 0.0003).

**Figure 5.**
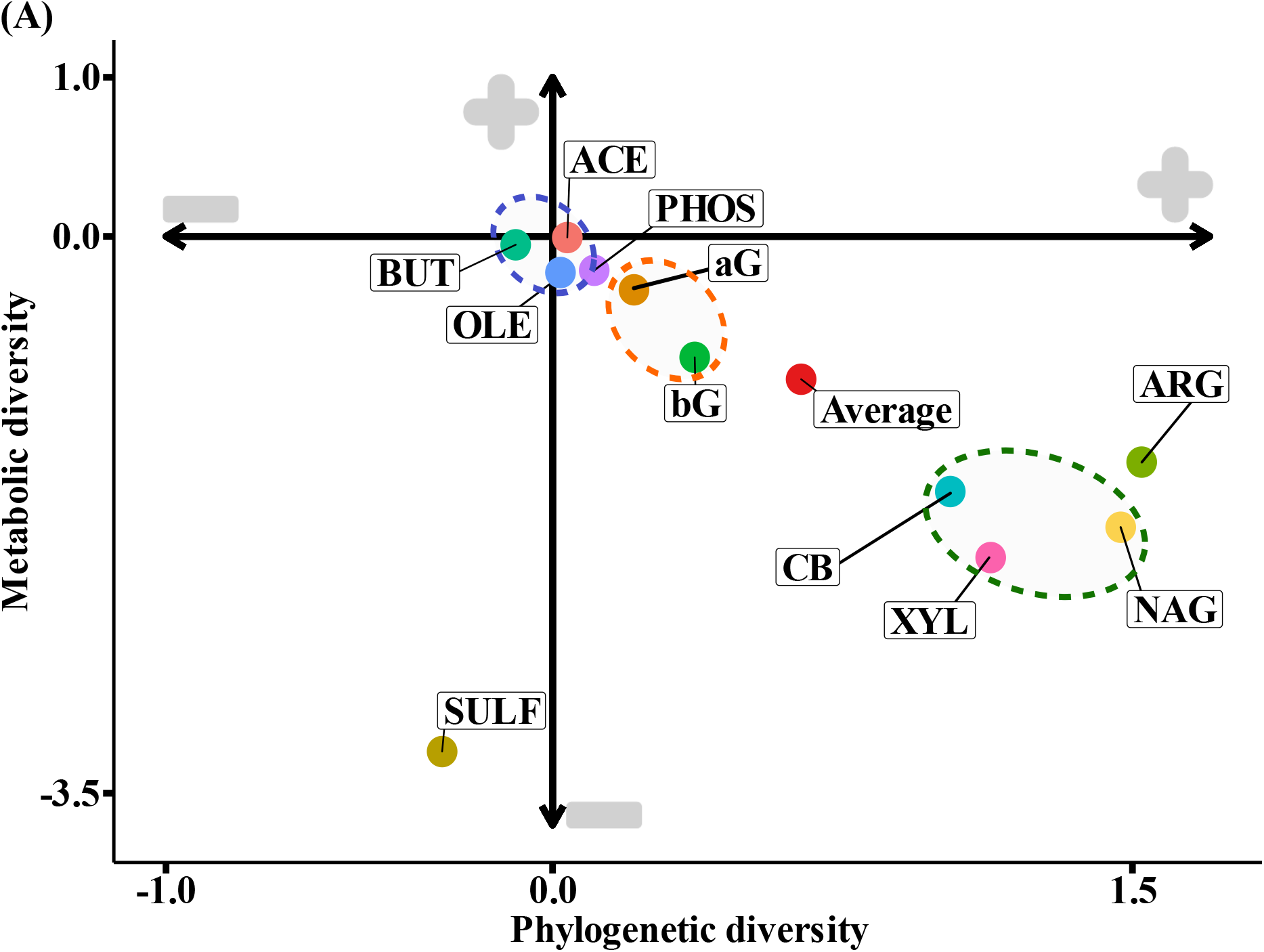
Substrate chemistry drives the strength and direction of phylogenetic and metabolic diversity treatment effects. (A) The linear regression model slopes for all 11 substrates and the average for all substrates (Appendix 1: Table S10) are plotted here as points. In (B) the average for all substrates, community EEA, is positively related with phylogenetic diversity (p < 0.001). In (C) the average for all substrates, community EEA, is negatively related with metabolic diversity (p < 0.001). Coloured points in (A) indicate different substrates and the average (red). Biochemically similar substrates are circled in dashed ellipses: blue for fatty acids, orange for monosaccharides and green for polysaccharides. Coloured squares in (B) and (C) indicate community groups: Psim-Msim in red, Psim-Mdis in green, Pdis-Msim in blue and Pdis-Mdis in purple.

### Effects of diversity in EEA changes over 24 h

We further investigated how the community EEA changed with increasing time after community mixing, by comparing EEA results from time points of 0 h, 6 h and 24 h (Figure 6). Significant positive changes ranged from 10 − 240% and negative changes ranged from −10 - −137%. Both Msim groups as well as Pdis-Mdis exhibited negative or non-significant changes over the 24 h. However, the Psim-Mdis group showed both significantly negative and positive changes between EEAs measured over 24 h after mixing. Changes in the Pdis communities’ composition within 24 h did not explain the observed activity changes; 16S rRNA gene amplicon sequencing results showed that all the introduced species appear to persist in the community cultures during 24 h post-mixing, with only close to ~1-fold change of relative abundance for most species and a 2-fold change for two species (Appendix S1: Figure S6). Thus, the observed differences in changes in EEA over time between diversity treatments were not the result of a shifting community composition, but of diversity treatment effects.

**Figure 6.**
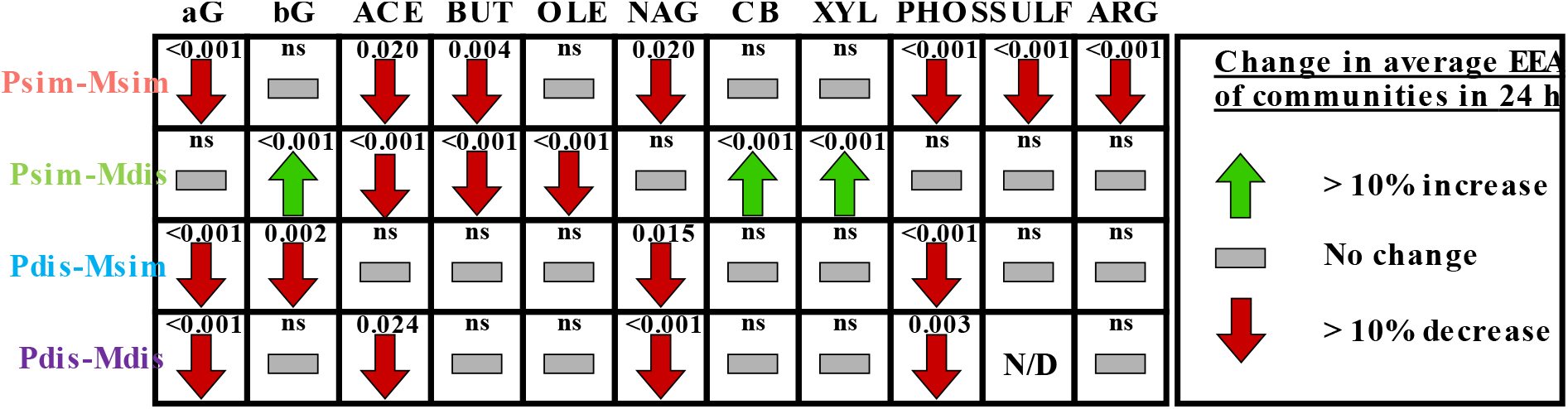
Temporal changes of EEA % proportional changes between communities and isolates from 0 h to 24 h after community mixing. Rows show community groups and columns show the substrate where the activity change was noted. The average changes between the timepoints are indicated by the colour of the arrows and their statistical significance (Student’s *t*-test, 2-tailed) showed on top of the arrows. P values below 0.001 are shown as <0.001 and values over 0.05 are marked as non-significant (*ns*).

### Substrate acclimation in EEA assays

Acclimation describes any phenotypic plasticity an organism can undergo to adjust to changes in its environment. Effects of substrate acclimation were observed directly by measuring the increase of EEA due to potential upregulation of exoenzyme production, during the incubation with a substrate. Here we show the results from the assay with aG, as an example. Comparing the timepoint of 0 h to the acclimated cultures timepoint of 4.5 h, we observed increases of the EEAs for both. However, these increases were much higher for communities, with ~5-fold increases on average, than for isolates, with average increases of ~1-fold (Figure 7). This was also reflected in the highest changes observed: For isolate activities in the aG substrate assay, 2.3 times increase, from 0.110 ± 0.003 to 0.255 ± 0.008 μM min^-1^, was the largest change observed. For community activities, increases up to 10 times, from 0.020 ± 0.001 to 0.199 ± 0.012 μM min^-1^ were observed, making community acclimation capacity almost 5 times higher compared to isolates. This increase in activity did not match the EEA changes observed for communities after 6 h of mixing without the addition of substrates. Since both isolate and community activities were normalised to biomass, the increased activity in communities could not be attributed to differences in cell density. In addition, the acclimation effect could not have been the result of species abundance changes since even the most active of the isolates only had ~2 times increase of EEA due to acclimation (0.225 ± 0.006 to 0.454 ± 0.003 μM min^-1^).

**Figure 7.**
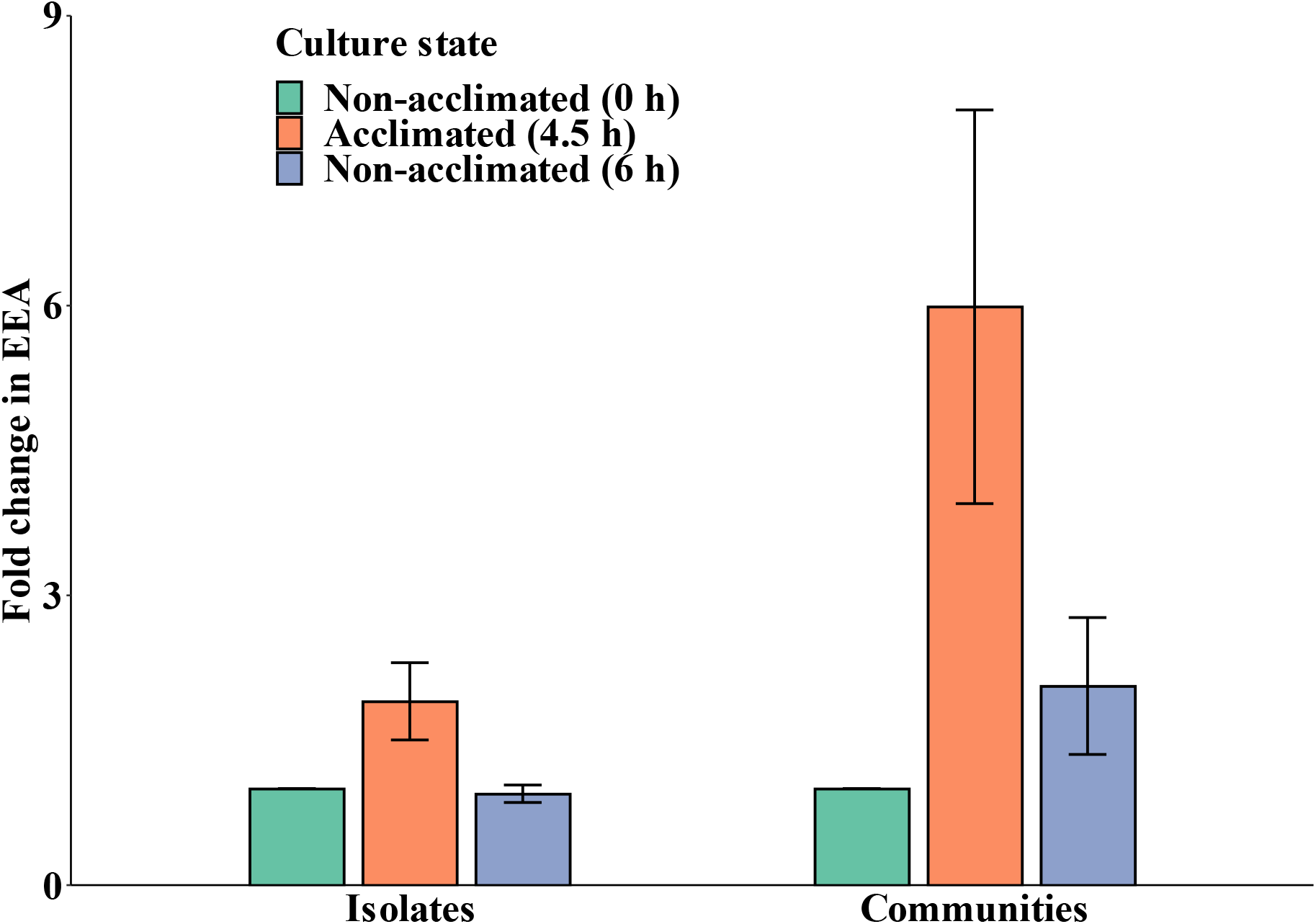
Substrate acclimation effect on isolates and communities. EEA of Psim-Mdis group isolates and communities on 4-MUB-α-D-glucopyranoside (aG) are shown as an example of the substrate acclimation effect. In green are shown the non-acclimated EEAs of cultures as a fold-change of 1, to which all other activities were normalised. In orange, the cultures received the substrate at the same time as the previous cultures and left to acclimate for 4.5 h. Lastly, the EEAs of cultures that had been mixed for 6 h but not left to acclimate, as shown in blue. Bars depict average fold-changes in the EEA of isolates and communities and error bars show the standard deviation (n = 5). Community EEAs showed a ~5-fold increase when acclimated with the substrate and isolates only showed a ~1-fold increase.

## Discussion

Using synthetic microbial communities, we showed that both phylogenetic and metabolic diversity influence community ecological functioning. Because both diversity aspects are so intricately linked, we used a multifactorial treatment experiment that allowed us to examine what the effects of each diversity were when controlling for the other. Since metabolic diversity has a direct impact on ecological functions, we expected that higher metabolic diversity would lead to higher functioning of bacterial communities. What we see is that the interaction between diversity treatments causes contrasting effects, which did not meet our expectations.

Metabolic diversity has either no effect, for Pdis communities, or a negative relationship, for Psim communities for both community TAC and community EEA % proportional changes (Figure 3B, Appendix S1: Figure S4B). In Pdis communities, differences in nutrient acquisition and interaction strategies of different species could be playing a role for the lack of effect of metabolic diversity. Such a mixture of species can engage in varying competitive or cooperative interactions (Gross 2008; Violle *et al*. 2011). This complementary set of community species interactions might be masking any observable influence of metabolic diversity (Maynard, Crowther and Bradford 2017). On the other hand, in Psim communities, related species are likely to be reacting similarly to each other and resource competition. However, increased conspecific competition can result in different outcomes. With lower metabolic diversity, upregulation of exoenzyme activity as a response to competition for resource exploitation, might be translating to increased community functioning. The mechanism behind this might be how the cost:benefit ratio of exoenzyme production manifests in low diversity communities if they are all producers and strong competitors (Allison *et al*. 2011). Consequently, higher metabolic diversity results in lower community functioning as species can diversify their resource preferences and therefore do not contribute to the same substrate degradation in the assays. Moreover, species of different genera might have distinct ways of interacting with species of their own genus, further influencing competitive outcomes (Mayfield and Levine 2010; Maynard, Crowther and Bradford 2017).

We also predicted that as phylogenetic diversity is based on a neutral genetic marker, it would have no effect on community ecological functioning. We found that for both TAC and EEA % changes, phylogenetic diversity has a positive relationship with the ecological functioning of Mdis communities and a negative relationship for Msim communities (Figure 3A; Appendix S1: Figure S4A). In the Mdis communities, organisms can engage in a variety of metabolic functions. This increases the capacity for different kinds of community species interactions (Weber and Strauss 2016), facilitated by phylogenetic diversity and coexistence. More diverse species interactions might translate into more cooperative communities, metabolic complementarity and increased overall functioning. Contrastingly, Msim communities rely on a limited number of metabolic functions. The limited metabolic functions do not allow for elaborate interaction strategies, such as cooperation, to be enacted. As such, any influence of phylogenetic diversity is severely decreased. In this setup, resource competition and antagonistic interactions with multiple species cause a negative relationship to emerge.

Lastly, we hypothesised that because Psim community species would have less diverse strategies in responding to situations of competition or cooperation, metabolic diversity would have a much stronger effect in their ecological functioning, compared to their Pdis community member counterparts. Indeed, this is true as we show that Psim communities had a strong, albeit negative relationship with EEA changes and TAC, while Pdis communities had none. Moreover, the consistency of results between the single and multifunctional measures of EEAs and TAC increases the confidence of the diversity relationships we have observed. This should not be surprising, as we already established a deep connection between phylogenetic and metabolic diversity, which would explain why neither of them is the sole driver of community ecological functions.

The contrasting diversity effects were also revealed when controlling for the substrates in the treatments. We identified as a general trend that phylogenetic diversity has a positive effect on EEA changes and metabolic diversity has a negative influence (Figure 5). In addition to that, we saw that the diversity effects were operating on a strength gradient, with groups of biochemically similar substrates appearing in distinct positions along that gradient (Figure 5). Such differences between substrates and groups of substrates can be related to inherent variations in exoenzyme production, maintenance and regulation and how different classes of enzymes behave to break down their substrates. For example, polysaccharides are targeted by hydrolytic exoenzymes that are produced on-demand (Warren 1996; Lynd *et al*. 2002). In contrast, enzymes involved in the degradation of fatty acids belonging to the ester group of substrates are part of bacterial core metabolism, and thus can be continuously produced. Our results show that stronger diversity effects are more important to ecosystem functions that rely on environmental factors, such as the temporal or spatial presence of plant-derived polysaccharides. Further adding to the complexity of experimental outcomes, time also influences different substrates and diversity treatments in different ways (Figure 6). Hence, not only the diversity of ecosystem functions tested (Huston 1997, 2000; Tilman 1997; Loreau 2000) but also their underlying biochemical mechanisms have a great effect on the outcomes of biodiversity research and the choice, effect and timing of different functions in experimental designs needs to considered.

To understand the very strong contrasting changes in EEAs between the Psim groups, their genus-specific capacity for co-existence with its related species is crucial. Psim-Msim communities contained *Flavobacteria* species which are usually non-motile organisms, prefer aquatic environments and have strong differences between species. The genus includes generalist species that can use a wide range of carbohydrates, to more specialist asaccharolytic species that prefer nitrogenous substrates (i.e. amino acids) (Bernardet and Bowman 2006). These physiological characteristics might allow the *Flavobacteria* communities to increase their EEA as different species could be utilising different carbon sources in their growth medium. Communities of these specialist *Flavobacteria* could be facilitating community species interactions that translate into increases in their community EEAs. In contrast, the *Pseudomonas* species in Psim-Mdis communities are generally motile-planktonic bacteria, ubiquitous in almost all oxic environments, including soil and water, due to their expansive metabolic repertoire, simple nutritional requirements and rapid growth rates (Moore *et al*. 2006). However, the reduced capacity of many *Pseudomonas* species to withstand starvation (Moore *et al*. 2006) might lead them to adapt more selfish nutrient acquisition strategies (Reintjes *et al*. 2017, 2019; Arnosti, Reintjes and Amann 2018). As generalist species, *Pseudomonas* might be more competitive, resulting in *Pseudomonas* communities with lower EEAs than their respective isolates. Thus, bacteria with a broad substrate range might not benefit when living in communities of the same genus but of dissimilar metabolism. Contrastingly, in the Pdis communities these genus-specific effects in nutrient acquisition strategies can be balanced due to the increased species diversity. Nevertheless, even competing generalists like *Pseudomonas* species will profit from species interactions when incubated for longer time in the presence of certain substrates. Using the Psim-Mdis communities to run EEA substrate acclimated vs. non-acclimated assays (Figure 7), we identified that the amplifying effect of substrate acclimation on the EEAs is much stronger for the communities than for the constituent isolates on substrate aG. In addition, since the increased EEAs are not present in the non-acclimated 6 h timepoint communities, we can conclude that the effect was indeed substrate acclimatisation of the bacteria and not an effect of mixing the community cultures. Consequently, the observed acclimation effect was the result of community species interactions that enhanced species’ ability to acclimate to the substrate. These observations link back to our first hypothesis, where we predict that the metabolic diversity of the group will enhance the functioning of communities. But oversimplifying the experimental design and just looking at one substrate or one group, would have let to an underestimation of the complexity of diversity-ecosystem functioning relationships, possibly explaining as to why we see studies showing opposite effects of biodiversity (Balvanera *et al*. 2006).

Altogether, our results concur with previous studies that focus on separating the different aspects of biodiversity and examining the relationship between biodiversity and ecosystem functioning (Griffiths *et al*. 2001; Balvanera *et al*. 2006; Zhang and Zhang 2006; Naeem *et al*. 2009; Jousset *et al*. 2011; Purschke *et al*. 2013; Thompson, Davies and Gonzalez 2015; Gamfeldt and Roger 2017). A meta-analysis demonstrated that biodiversity effects depend on the organisation level: with positive effects at the community level, positive but weaker effects at the ecosystem level and negative effects at the population level (Balvanera *et al*. 2006). Experiments done by Griffiths *et al*. (Griffiths *et al*. 2001) recognised no consistent effect from biodiversity on soil functions. Thus, the need for more research including the effects of the environmental context, like nutrient availability, on the relationship between biodiversity and ecosystem functioning was recognized (Zhang and Zhang 2006). Our study fills this gap by focussing not only on the phylogenetic and metabolic diversity of organisms, but also measuring a broad diversity of activities controlled by different underlying mechanisms of enzyme regulation. Such a complex approach is required to tackle the complex dimensionality of biodiversity research. We show that biodiversity had a significant impact on most EEAs, but the strength of the effect depended on the substrate context. This might explain why the relationship between biodiversity and community ecosystem functioning cannot always be predicted to be positive as previously observed (Aarssen 1997; Tilman 1997; Huston 2000; Carroll, Cardinale and Nisbet 2011). Natural, larger scale experiments struggle with introducing different levels of complexity in their experimental design (Cairns *et al*. 2018). Looking forward, we should aim to understand how diversity effects can mechanistically influence ecosystems, synthetic and natural (Fox and Harpole 2008; Naeem *et al*. 2009; Turnbull *et al*. 2013), and not take the effects as granted (Shade 2017).

## Acknowledgements

We gratefully acknowledge the support of the German Centre for Integrative Biodiversity Research (iDiv) Halle-Jena-Leipzig funded by the Deutsche Forschungsgemeinschaft (DFG, German Research Foundation) (DFG FZT 118, 202548816). This project has been conducted in the framework of the iDiv-Flexpool – the internal funding mechanism of iDiv. Martin Taubert was funded by the DFG under Germany’s Excellence Strategy – EXC 2051 – Project-ID 390713860. We would like to express our gratitude to Patricia Geesink for the bacterial isolation work under the framework of the CRC AquaDiva of the Friedrich Schiller University Jena, funded by the DFG (DFG SFB 1076). Additionally, we would like to thank Will A. Overholt and Lijuan Yan for their invaluable statistical analysis input.

## Author Contributions

KK and SWH secured funding for the project. CX, MT, and KK designed the experiments. CX performed the experiments and carried out all assays and molecular analyses; the data analysis was performed by CX closely supported by MT. Statistical analysis was performed by CX supported by MT and SWH. All co-authors assisted in interpreting the results; CX initiated the manuscript writing, which was finalized with substantial contributions from MT, KK and SWH.

