## Appendix S1 for "Phylogenetic and metabolic diversity have contrasting effects on the ecological functioning of bacterial communities"

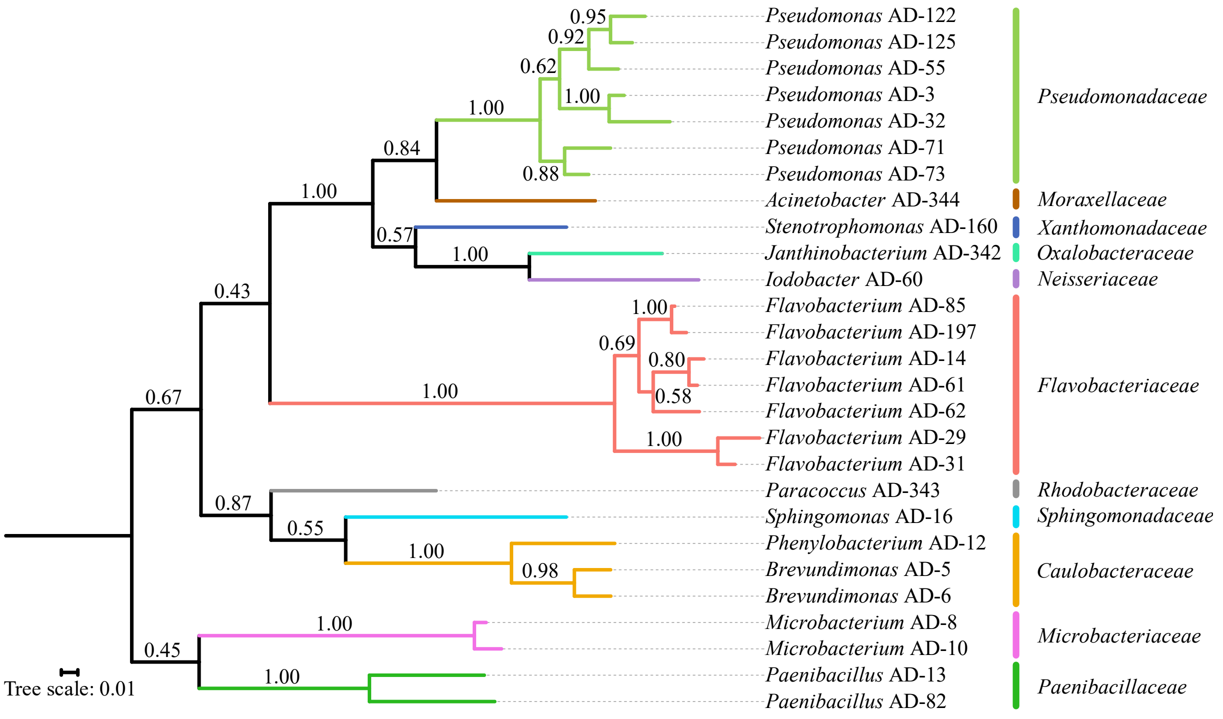

**Figure S1: Bacterial isolate phylogenetic tree.** The dendrogram was calculated using the maximum likelihood method and Tamura-Nei model in MEGAX (Kumar et al. 2018) based on full-length 16S rRNA gene sequences (universal primers 27F and 1492R) of groundwater bacterial isolates (500 bootstrap replicates). Node labels depict the bootstrap values and the scale bar shows the divergence time for each clade. The different colours denote taxonomic families.

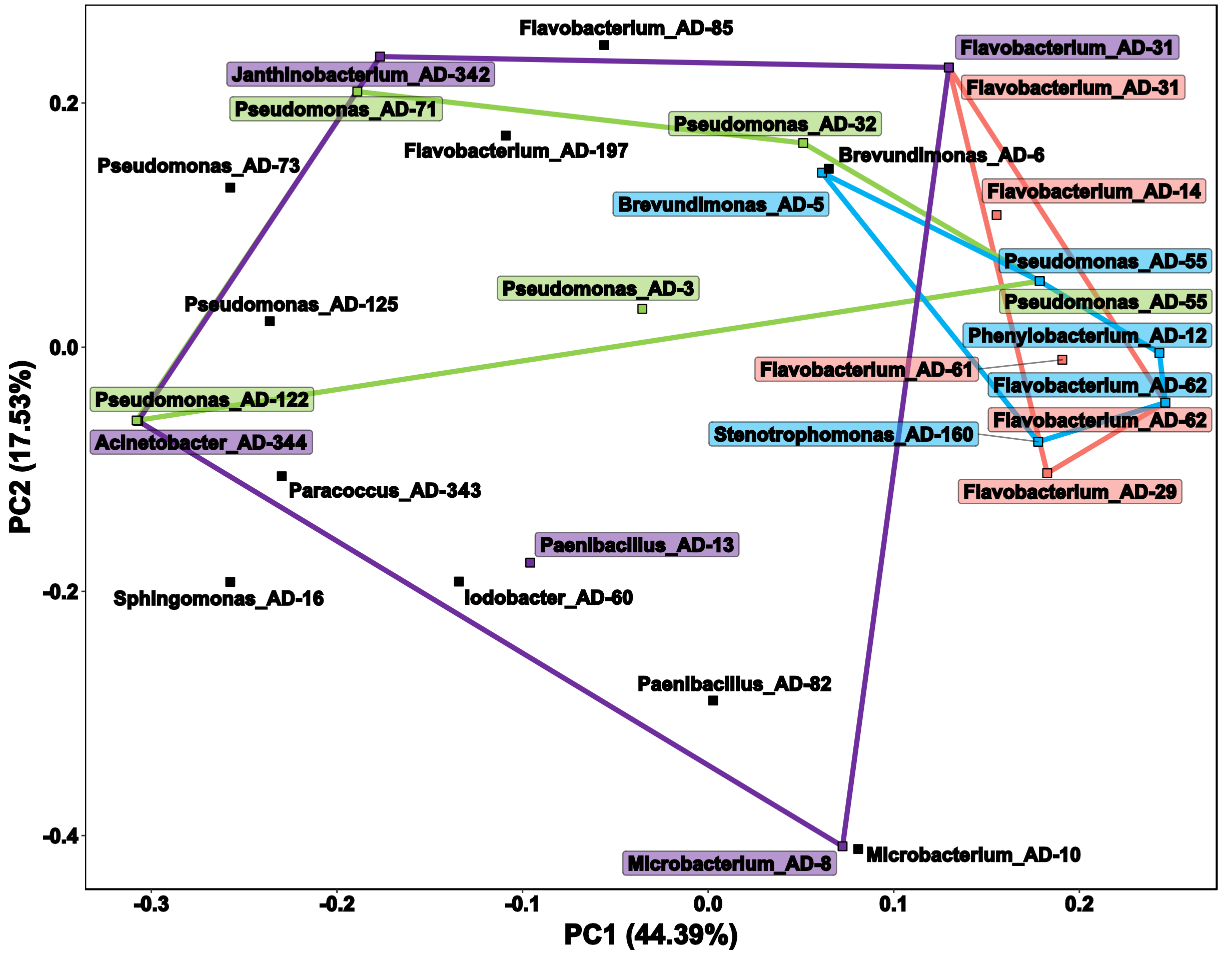

**Figure S2: PCA based on EEAs of bacterial isolates**. The analysis was performed in R (v.3.6.1) (R Core Team 2019). Colours indicate the groups the isolates were assigned to, with red for Psim-Msim group, green for Psim-Mdis group, blue for Pdis-Msim group and purple for Pdis-Mdis group. Duplicate isolate labels indicate isolates that were assigned to more than one community group.

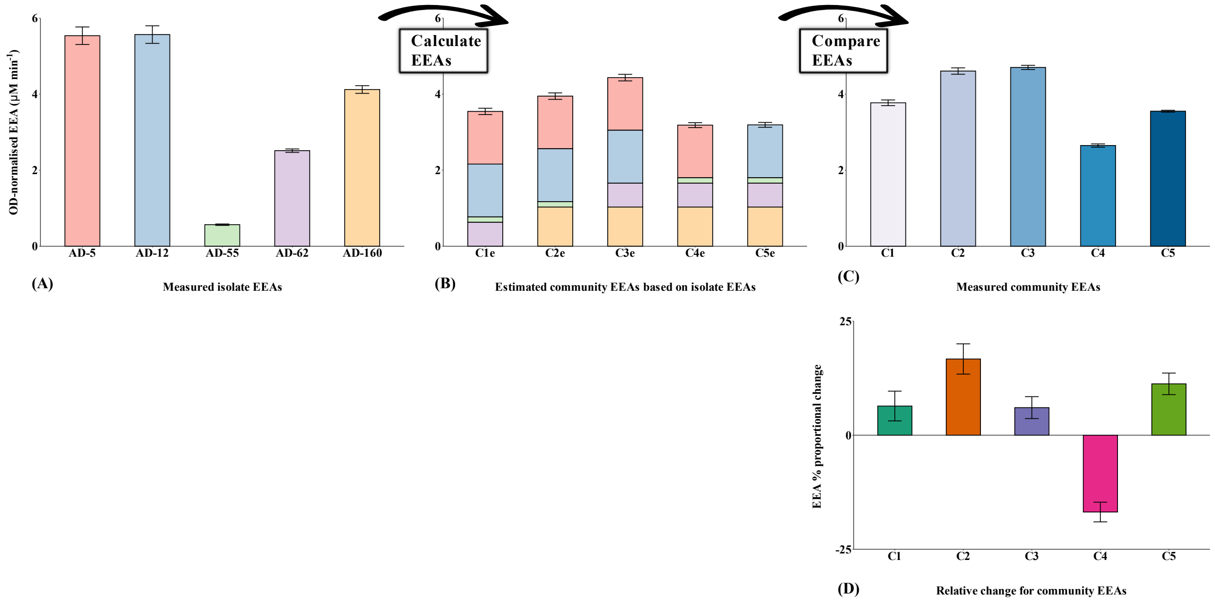

**Figure S3: Data transformation for the comparison of community exoenzyme activities to the activity of their constituent isolates.** (A) An example of the EEAs of five isolates belonging to the same community group (here as isolates of the Pdis-Msim group for the ACE assay). Their activities are used to calculate (B) the expected activity of their community cultures, as they are mixed at similar biomass levels (labelled as C1-C5$_{\text{expected (e)}}$. (C) The measured activity from the experimental communities is compared to the expected activities and given as (D) proportional change in EEA. The proportional change data is used in subsequent analyses and comparisons.

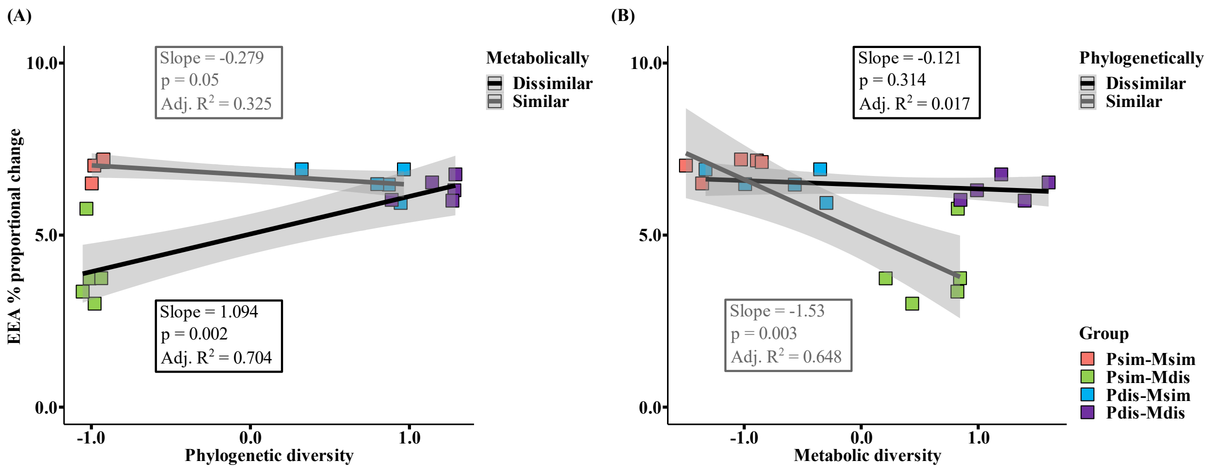

**Figure S4: Effects of phylogenetic diversity (A) and metabolic diversity (B) on the EEA proportional changes of communities.** Black regression lines show the effect of each treatment when controlling for the dissimilar groups of the second treatment. Grey regression lines show the effect of each treatment when controlling for the similar groups of the second treatment. Statistical significance of nested ANOVAs for each linear regression is shown in each plot, with the respective coloured text (grey or black). Coloured squares in (B) and (C) indicate community groups: Psim-Msim in red, Psim-Mdis in green, Pdis-Msim in blue and Pdis-Mdis in purple. The y axes have been log2 transformed and the x axes have been z-scale transformed.

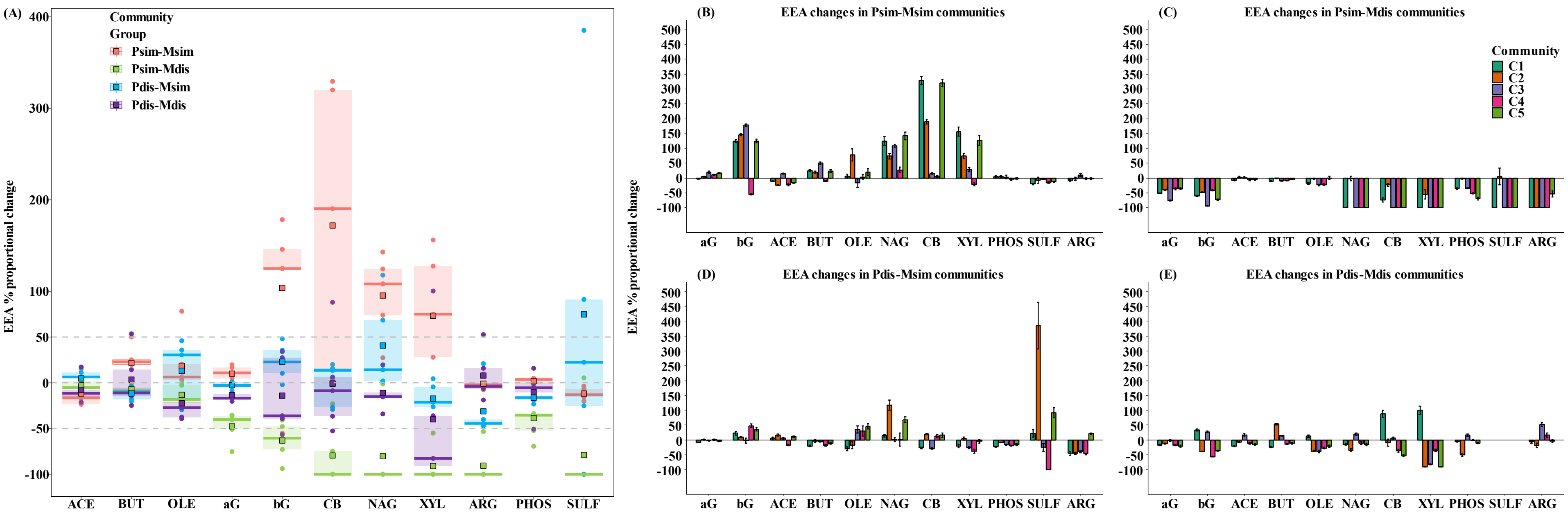

**Figure S5: EEA proportional changes for communities compared to their constituent isolates.** (A) The EEAs changes for each of the four community groups are shown, with in their mean activity for each substrate in square points. Coloured solid lines and shaded areas respectively show the median and quartiles of the data for each community within the group. Points show proportional change values for each community within the group. Pdis groups (blue and purple) EEA changes remain within the ±50% change thresholds, indicated by the grey dashed lines. The bar plots show EEA changes for each community culture individually: in (B) for the Psim-Msim group, (C) Psim-Mdis, (D) Pdis-Msim, (E) Pdis-Mdis.

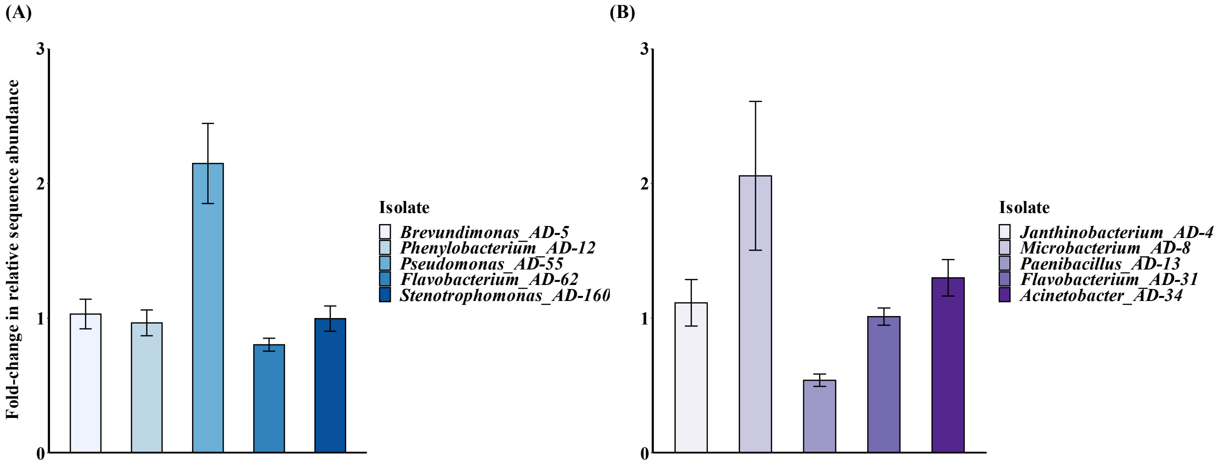

**Figure S6: Changes in microbial community composition over time**. The average changes in species abundance for all communities of the (A) Pdis-Msim group and (B) Pdis-Mdis group are shown as fold change from 0 h to 24 h after mixing them. Data were derived from Illumina MiSeq 16S amplicon sequencing of the V3-V4 region of the 16S rRNA gene. Bars show the mean fold-change in the relative abundance of each species for the Pdis groups and the error bars show the standard deviation.

**Table S1. Taxonomy and Genbank accession information of bacterial isolates used in this study.** Isolates with strain information "hainich_*" have been previously published (Geesink et al., 2018).

| Isolate Genus_ID | Strain | Genbank Accession No. | NCBI Taxonomy |
|---|---|---|---|
| Pseudomonas_AD-3 | hainich_003 | MG980419 | Proteobacteria; Gammaproteobacteria; Pseudomonadales; Pseudomonadaceae |
| Brevundimonas_AD-5 | hainich_005 | MG980421 | Proteobacteria; Alphaproteobacteria; Caulobacterales; Caulobacteraceae |
| Brevundimonas_AD-6 | hainich_006 | MG980422 | Proteobacteria; Alphaproteobacteria; Caulobacterales; Caulobacteraceae |
| Microbacterium_AD-8 | hainich_008 | MG980423 | Actinobacteria; Actinobacteria; Micrococcales; Microbacteriaceae |
| Microbacterium_AD-10 | hainich_010 | MG980425 | Actinobacteria; Actinobacteria; Micrococcales; Microbacteriaceae |
| Phenylobacterium_AD-12 | hainich_012 | MG980426 | Proteobacteria; Alphaproteobacteria; Caulobacterales; Caulobacteraceae |
| Paenibacillus_AD-13 | hainich_013 | MG980427 | Firmicutes; Bacilli; Bacillales; Paenibacillaceae |
| Flavobacterium_AD-14 | hainich_014 | MG980428 | Bacteroidetes; Bacteroidia; Flavobacteriales; Flavobacteriaceae |
| Sphingomonas_AD-16 | hainich_016 | MG980430 | Proteobacteria; Alphaproteobacteria; Sphingomonadales; Sphingomonadaceae |
| Flavobacterium_AD-29 | hainich_029 | MG980438 | Bacteroidetes; Bacteroidia; Flavobacteriales; Flavobacteriaceae |
| Flavobacterium_AD-31 | hainich_031 | MG980439 | Bacteroidetes; Bacteroidia; Flavobacteriales; Flavobacteriaceae |
| Pseudomonas_AD-32 | hainich_032 | MG980440 | Proteobacteria; Gammaproteobacteria; Pseudomonadales; Pseudomonadaceae |
| Pseudomonas_AD-55 | hainich_055 | MG980457 | Proteobacteria; Gammaproteobacteria; Pseudomonadales; Pseudomonadaceae |
| Iodobacter_AD-60 | hainich_060 | MG980462 | Proteobacteria; Betaproteobacteria; Neisseriaceae |
| Flavobacterium_AD-61 | hainich_061 | MG980463 | Bacteroidetes; Bacteroidia; Flavobacteriales; Flavobacteriaceae |
| Flavobacterium_AD-62 | hainich_062 | MG980464 | Bacteroidetes; Bacteroidia; Flavobacteriales; Flavobacteriaceae |
| Pseudomonas_AD-71 | hainich_071 | MG980471 | Proteobacteria; Gammaproteobacteria; Pseudomonadales; Pseudomonadaceae |
| Pseudomonas_AD-73 | hainich_073 | MG980473 | Proteobacteria; Gammaproteobacteria; Pseudomonadales; Pseudomonadaceae |
| Paenibacillus_AD-82 | hainich_082 | MG980481 | Firmicutes; Bacilli; Bacillales; Paenibacillaceae |
| Flavobacterium_AD-85 | AD-85 | MN547321 | Bacteroidetes; Bacteroidia; Flavobacteriales; Flavobacteriaceae |
| Pseudomonas_AD-122 | AD-122 | MN547322 | Proteobacteria; Gammaproteobacteria; Pseudomonadales; Pseudomonadaceae |
| Pseudomonas_AD-125 | AD-125 | MN547323 | Proteobacteria; Gammaproteobacteria; Pseudomonadales; Pseudomonadaceae |
| Stenotrophomonas_AD-160 | hainich_160 | MG980541 | Proteobacteria; Gammaproteobacteria; Xanthomonadales; Xanthomonadaceae |
| Flavobacterium_AD-197 | hainich_197 | MG980562 | Bacteroidetes; Bacteroidia; Flavobacteriales; Flavobacteriaceae |
| Janthinobacterium_AD-342 | AD-342 | MN547318 | Proteobacteria; Betaproteobacteria; Burkholderiales; Oxalobacteraceae |
| Paracoccus_AD-343 | AD-343 | MN547319 | Proteobacteria; Alphaproteobacteria; Rhodobacterales; Rhodobacteraceae |
| Acinetobacter_AD-344 | AD-344 | MN547320 | Proteobacteria; Gammaproteobacteria; Pseudomonadales; Moraxellaceae |

**Table S2. Catabolic exoenzyme-targeting substrates with a fluorescent moiety**; Methylumbelliferyl (MUB) or Amino-4-ethylcoumarin (AMC) and standards used for each. In total, eleven substrates were used to characterize the exoenzymatic activity of bacterial isolates and synthetic communities.

| Enzyme Target | Category | Enzyme(s) Action | Fluorescent Substrate | Solvent |
|---|---|---|---|---|
| Cellulase | β-Glucanases | Hydrolysis of 1,4-beta-D-glucosidic linkages in cellulose, lichenin and cereal β-D-glucans | 4-MUB-β-D-cellobioside | $H_2O$ |
| α-Glucosidase | Glycosidases | Degrade complex sugars (starch, glycogen) | 4-MUB-α-D-glucopyranoside | DMSO |
| β-Glucosidase | Glycosidases | Degrade complex sugars (starch, glycogen) | 4-MUB-β-D-glucopyranoside | $H_2O$ |
| Leucine aminopeptidase | Exopeptidases | Degrade Proteins | Z-L-Arginine-7-AMC HCL | DMSO |
| N-acetyl-β-Glucosaminidase | Glycosidases | Chitin hydrolysis (glycosidic bonds break) | 4-MUB-N-acetyl-β-D-glucosaminide | DMSO |
| Phosphatase | Esterases | Phosphorus mineralization from phosphate esters | 4-MUB phosphate | $H_2O$ |
| β-D-xylopyranosidase | Glycosidases | Degrade Hemicelluloses | 4-MUB-β-D-xylopyranoside | $H_2O$ |
| Butyrate Kinase | Transferase | Degrade plant waxes | 4-MUB-butyrate | DMSO |
| Lipases | Lipases | Fatty acids hydrolysis | 4-MUB oleate | DMSO |
| Sulfatase | Esterases | Sulfate esters hydrolysis | 4-MUB sulfate potassium salt | $H_2O$ |
| Acetate Kinase | Transferase | Aerobic respiration cycle | 4-MUB acetate | DMSO |
| AMC standard | Standard | - | 7-Amino-4-methylcoumarin | EtOH |
| MUB standard | Standard | - | 4-Methylumbelliferone | DMSO |

**Table S3. Isolate species composition for each community of each group and calculated phylogenetic and metabolic diversity for each community and for the whole group.** Shaded boxes (grey) indicate which bacterial isolates were included in each community. Reported phylogenetic diversity values are estimates of average evolutionary divergence over sequence pairs within groups. Reported metabolic diversity values stem from the Euclidean distance matrix used for the hierarchical clustering and quantification of the total EEAs for each isolate. Significant differences in the mean phylogenetic diversity were observed from groups Psim-Msim (0.072 ± 0.004) and Psim-Mdis (0.066 ± 0.005) to groups Pdis-Msim (of 0.293 ± 0.030) and Pdis-Mdis (0.343 ± 0.020) (paired sample *t* -test, t value = −19.37, $p < 0.0001$). Metabolic diversity differed significantly as well, from groups Psim-Msim (7.970 ± 0.823) and Pdis-Msim (9.300 ± 1.260) to groups Psim-Mdis (13.550 ± 0.825) and Pdis-Mdis (15.395 ± 0.864) (paired sample *t* -test, t value = −10.44, $p < 0.0001$).

| Community | Isolate 1 | Isolate 2 | Isolate 3 | Isolate 4 | Isolate 5 | Phylogenetic Diversity | Metabolic Diversity |
|---|---|---|---|---|---|---|---|
| **Psim-Msim** | ***Flavobacterium* AD-14** | ***Flavobacterium* AD-29** | ***Flavobacterium* AD-31** | ***Flavobacterium* AD-61** | ***Flavobacterium* AD-62** | | |
| **C1** | | | | | | 0.075 | 8.714 |
| **C2** | | | | | | 0.075 | 8.85 |
| **C3** | | | | | | 0.068 | 6.783 |
| **C4** | | | | | | 0.066 | 7.224 |
| **C5** | | | | | | 0.076 | 8.282 |
| | | | | | **Average** | **0.072 ± 0.004** | **7.970 ± 0.823** |
| **Psim-Mdis** | ***Pseudomonas* AD-3** | ***Pseudomonas* AD-32** | ***Pseudomonas* AD-55** | ***Pseudomonas* AD-71** | ***Pseudomonas* AD-122** | | |
| **C1** | | | | | | 0.065 | 12.218 |
| **C2** | | | | | | 0.062 | 14.182 |
| **C3** | | | | | | 0.069 | 12.941 |
| **C4** | | | | | | 0.059 | 14.169 |
| **C5** | | | | | | 0.074 | 14.242 |
| | | | | | **Average** | **0.066 ± 0.005** | **13.550 ± 0.825** |
| **Pdis-Msim** | ***Brevundimonas* AD-5** | ***Phenylobacterium* AD-12** | ***Pseudomonas* AD-55** | ***Flavobacterium* AD-62** | ***Stenotrophomonas* AD-160** | | |
| **C1** | | | | | | 0.305 | 9.746 |
| **C2** | | | | | | 0.235 | 10.435 |
| **C3** | | | | | | 0.295 | 8.395 |
| **C4** | | | | | | 0.314 | 10.605 |
| **C5** | | | | | | 0.317 | 7.317 |
| | | | | | **Average** | **0.293 ± 0.030** | **9.300 ± 1.261** |

| Pdis-Mdis | *Janthinobacterium* AD-342 | *Microbacterium* AD-8 | *Paenibacillus* AD-13 | *Flavobacterium* AD-31 | *Acinetobacter* AD-344 | | |
|---|---|---|---|---|---|---|---|
| **C1** | | | | | | 0.358 | 15.363 |
| **C2** | | | | | | 0.307 | 14.252 |
| **C3** | | | | | | 0.339 | 16.652 |
| **C4** | | | | | | 0.357 | 14.703 |
| **C5** | | | | | | 0.356 | 16.001 |
| | | | | | **Average** | **0.343 ± 0.020** | **15.395 ± 0.864** |

**Table S4. Linear regression models for the effects of phylogenetic and metabolicl diversity on the TAC of bacterial communities.** The effects of phylogenetic diversity were examined on the two subgroups of the metabolic diversity treatments, metabolically similar (Msim) and metabolically dissimilar (Mdis) communities. Likewise, the treatments for phylogenetic diversity (Psim, Pdis) were controlled when studying the effects of metabolic diversity. Significant effects are shown with their p values in **bold font**.

| | **Phylogenetic diversity** | | | | | **Metabolic diversity** | | | | | |
|---|---|---|---|---|---|---|---|---|---|---|---|
| ***Controlled treatment*** | ***Estimate*** | ***± S.E.*** | ***t*** | ***p*** | ***df*** | ***Estimate*** | ***± S.E.*** | ***t*** | ***p*** | ***df*** | $R^2$ / $R^2$ ***adjusted*** |
| **TAC of Msim** | -0.305 | 0.102 | -2.982 | **0.018** | 10 | | | | | | 0.526 / 0.467 |
| **TAC of Mdis** | 0.313 | 0.105 | 2.981 | **0.018** | 10 | | | | | | 0.526 / 0.467 |
| **TAC of Psim** | | | | | | -0.611 | 0.151 | -4.054 | **0.004** | 10 | 0.673 / 0.632 |
| **TAC of Pdis** | | | | | | 0.002 | 0.115 | 0.018 | 0.986 | 10 | 0.000 / -0.125 |

**Table S5. Linear regression models for the effects of phylogenetic and metabolic diversity on the EEA proportional changes of bacterial communities.** The effects of phylogenetic diversity were examined on the two subgroups of the metabolic diversity treatments, metabolically similar (Msim) and metabolically dissimilar (Mdis) communities. Likewise, the treatments for phylogenetic diversity (Psim, Pdis) were controlled when studying the effects of metabolic diversity. Significant effects are shown with their p values in **bold font.**

| | **Phylogenetic diversity** | | | | | **Metabolic diversity** | | | | | |
|---|---|---|---|---|---|---|---|---|---|---|---|
| ***Controlled treatment*** | ***Estimate*** | ***± S.E.*** | ***t*** | ***p*** | ***df*** | ***Estimate*** | ***± S.E.*** | ***t*** | ***p*** | ***df*** | ***$R^2$ / $R^2$ adjusted*** |
| **EEAs of Msim** | -0.279 | 0.121 | -2.310 | **0.050** | 10 | | | | | | 0.400 / 0.325 |
| **EEAs of Mdis** | 1.094 | 0.231 | 4.738 | **0.002** | 10 | | | | | | 0.737 / 0.704 |
| **EEAs of Psim** | | | | | | -1.535 | 0.366 | -4.190 | **0.003** | 10 | 0.687 / 0.648 |
| **EEAs of Pdis** | | | | | | -0.121 | 0.113 | -1.074 | 0.314 | 10 | 0.126 / 0.017 |

**Table S6.** Psim-Msim EEAs for the 0h timepoint. Estimated community EEAs were calculated using the average EEAs of the constituent isolates. % proportional change of community EEAs was calculated as:
Community % change = (Measured community EEA - Estimated community EEA) / Estimated community EEA * 100

| | Cleaved substrate concentration - Measured (µM min$^{-1}$) | | | | | | | | | | |
|---|---|---|---|---|---|---|---|---|---|---|---|
| **Isolate** | **aG** | **bG** | **ACE** | **BUT** | **OLE** | **NAG** | **CB** | **XYL** | **PHOS** | **SULF** | **ARG** |
| **AD-14** | 0.10268 | 0.20568 | 1.16456 | 2.65751 | 0.00586 | 0.00686 | 0.04855 | 0.00552 | 0 | 0 | 0.00428 |
| **AD-29** | 0.98082 | 0.97762 | 5.01344 | 4.87659 | 0 | 0.0148 | 0.02821 | 0.00683 | 0.02129 | 0 | 0.03265 |
| **AD-31** | 0.24962 | 0.08648 | 6.78264 | 5.91827 | 0.00817 | 0.09093 | 0.07615 | 0 | 2.84093 | 0 | 0.07795 |
| **AD-61** | 0.28702 | 0.04285 | 2.23053 | 3.62031 | 0.01775 | 0.00426 | 0.00201 | 0.00417 | 0.00645 | 0.03349 | 0.04461 |
| **AD-62** | 1.14769 | 0.41201 | 2.91214 | 1.83133 | 0.00569 | 0.02052 | 0.07025 | 0.0104 | 0.00352 | 0.01002 | 0.02692 |
| **Community** | | | | | | | | | | | |
| **C1** | 0.39492 | 0.7379 | 3.40507 | 5.34109 | 0.00845 | 0.06555 | 0.16635 | 0.01058 | 0.75231 | 0.0067 | 0.03677 |
| **C2** | 0.65149 | 1.03391 | 3.03192 | 4.58637 | 0.00878 | 0.05789 | 0.16194 | 0.00995 | 0.74916 | 0.00234 | 0.03464 |
| **C3** | 0.75348 | 1.13991 | 3.25213 | 4.86995 | 0.00612 | 0.02416 | 0.043 | 0.00861 | 0.00809 | 0.01047 | 0.02967 |
| **C4** | 0.49532 | 0.08374 | 2.52334 | 3.14069 | 0.0097 | 0.03903 | 0.05128 | 0.00398 | 0.68973 | 0.00919 | 0.03756 |
| **C5** | 0.77752 | 0.85304 | 3.53761 | 4.99436 | 0.00952 | 0.07924 | 0.18552 | 0.01217 | 0.70788 | 0.00946 | 0.04447 |

| | Cleaved substrate concentration - Estimated community EEAs (µM min$^{-1}$) | | | | | | | | | | |
|---|---|---|---|---|---|---|---|---|---|---|---|
| **Community** | **aG** | **bG** | **ACE** | **BUT** | **OLE** | **NAG** | **CB** | **XYL** | **PHOS** | **SULF** | **ARG** |
| **C1e** | 0.40504 | 0.32816 | 3.79779 | 4.26817 | 0.00794 | 0.02921 | 0.03873 | 0.00413 | 0.71717 | 0.00837 | 0.03987 |
| **C2e** | 0.62021 | 0.42045 | 3.96819 | 3.82092 | 0.00493 | 0.03328 | 0.05579 | 0.00569 | 0.71643 | 0.00251 | 0.03545 |
| **C3e** | 0.62955 | 0.40954 | 2.83017 | 3.24643 | 0.00733 | 0.01161 | 0.03726 | 0.00673 | 0.00781 | 0.01088 | 0.02711 |
| **C4e** | 0.44675 | 0.18676 | 3.27247 | 3.50685 | 0.00937 | 0.03064 | 0.04924 | 0.00502 | 0.71272 | 0.01088 | 0.03844 |
| **C5e** | 0.66629 | 0.37974 | 4.23469 | 4.06162 | 0.0079 | 0.03263 | 0.04416 | 0.00535 | 0.71805 | 0.01088 | 0.04553 |

| | % proportional change in community EEAs | | | | | | | | | | |
|---|---|---|---|---|---|---|---|---|---|---|---|
| **Community** | **aG** | **bG** | **ACE** | **BUT** | **OLE** | **NAG** | **CB** | **XYL** | **PHOS** | **SULF** | **ARG** |
| **C1** | -2.49906 | 124.863 | -10.3409 | 25.1378 | 6.34921 | 124.407 | 329.515 | 156.142 | 4.90006 | -20.0182 | -7.77795 |
| **C2** | 5.0443 | 145.908 | -23.5944 | 20.033 | 78.2012 | 73.9445 | 190.261 | 74.895 | 4.5682 | -6.6378 | -2.29162 |
| **C3** | 19.6841 | 178.339 | 14.9096 | 50.0093 | -16.4629 | 108.076 | 15.4287 | 27.9671 | 3.56618 | -3.7841 | 9.43626 |
| **C4** | 10.8696 | -55.1585 | -22.8918 | -10.4414 | 3.50118 | 27.3814 | 4.14309 | -20.7248 | -3.22598 | -15.5437 | -2.27748 |
| **C5** | 16.6941 | 124.637 | -16.4611 | 22.9647 | 20.4183 | 142.871 | 320.141 | 127.534 | -1.41624 | -13.0671 | -2.33179 |

**Table S7.** Psim-Mdis EEAs for the 0h timepoint. Estimated community EEAs were calculated using the average EEAs of the constituent isolates. % proportional change of community EEAs was calculated as:
Community % change = (Measured community EEA - Estimated community EEA) / Estimated community EEA * 100

**Cleaved substrate concentration - Measured (µM min$^{-1}$)**

| Isolate | aG | bG | ACE | BUT | OLE | NAG | CB | XYL | PHOS | SULF | ARG |
|---|---|---|---|---|---|---|---|---|---|---|---|
| **AD-14** | 0.22536 | 0.07358 | 0.59874 | 4.36289 | 0.05647 | 0 | 0.00151 | 0.00177 | 0.06931 | 0.00347 | 0 |
| **AD-29** | 0.01522 | 0.01598 | 0.85944 | 2.77345 | 0.04207 | 0.00646 | 0.01802 | 0 | 0.06491 | 0.00232 | 0.00036 |
| **AD-31** | 0.10961 | 0.2055 | 1.28393 | 5.22384 | 0.04645 | 0.00868 | 0.02729 | 0.01053 | 0.00749 | 0 | 0.00047 |
| **AD-61** | 0 | 0 | 0.11819 | 0.2168 | 0.06765 | 0 | 0 | 0 | 0 | 0 | 0 |
| **AD-62** | 0 | 0 | 0.61939 | 4.11769 | 0.0481 | 0 | 0 | 0 | 0 | 0 | 0 |
| **Community** | | | | | | | | | | | |
| **C1** | 0.04287 | 0.02912 | 0.66845 | 2.81812 | 0.04343 | 0 | 0.00297 | 0 | 0.02287 | 0 | 0 |
| **C2** | 0.05228 | 0.03847 | 0.87184 | 4.08351 | 0.04696 | 0.00373 | 0.00901 | 0.00138 | 0.03466 | 0.00153 | 0 |
| **C3** | 0.01485 | 0.00139 | 0.55665 | 2.60533 | 0.04106 | 0 | 0 | 0 | 0.02222 | 0 | 0 |
| **C4** | 0.05335 | 0.04133 | 0.6156 | 3.17173 | 0.04273 | 0 | 0 | 0 | 0.00915 | 0 | 0 |
| **C5** | 0.02013 | 0.01501 | 0.6833 | 2.92107 | 0.05104 | 0 | 0 | 0 | 0.00554 | 0 | 9.7E-05 |

**Cleaved substrate concentration - Estimated community EEAs (µM min$^{-1}$)**

| Community | aG | bG | ACE | BUT | OLE | NAG | CB | XYL | PHOS | SULF | ARG |
|---|---|---|---|---|---|---|---|---|---|---|---|
| **C1e** | 0.08755 | 0.07377 | 0.71508 | 3.14425 | 0.05316 | 0.00379 | 0.0117 | 0.00307 | 0.03543 | 0.00145 | 0.00021 |
| **C2e** | 0.08755 | 0.07377 | 0.84037 | 4.11947 | 0.04827 | 0.00379 | 0.0117 | 0.00307 | 0.03543 | 0.00145 | 0.00021 |
| **C3e** | 0.06014 | 0.02239 | 0.54894 | 2.86771 | 0.05357 | 0.00162 | 0.00488 | 0.00044 | 0.03356 | 0.00145 | 9.1E-05 |
| **C4e** | 0.08374 | 0.06977 | 0.65506 | 3.48031 | 0.05467 | 0.00217 | 0.0072 | 0.00307 | 0.0192 | 0.00087 | 0.00012 |
| **C5e** | 0.03121 | 0.05537 | 0.72024 | 3.08294 | 0.05107 | 0.00379 | 0.01133 | 0.00263 | 0.0181 | 0.00058 | 0.00021 |

**% proportional change in community EEAs**

| Community | aG | bG | ACE | BUT | OLE | NAG | CB | XYL | PHOS | SULF | ARG |
|---|---|---|---|---|---|---|---|---|---|---|---|
| **C1** | -51.0336 | -60.5193 | -6.51994 | -10.372 | -18.3065 | -100 | -74.6162 | -100 | -35.4515 | -100 | -100 |
| **C2** | -40.2774 | -47.8547 | 3.74363 | -0.87275 | -2.72137 | -1.37988 | -22.988 | -54.9791 | -2.17372 | 5.39996 | -100 |
| **C3** | -75.3049 | -93.7859 | 1.40391 | -9.14933 | -23.362 | -100 | -100 | -100 | -33.7809 | -100 | -100 |
| **C4** | -36.292 | -40.7612 | -6.02379 | -8.86646 | -21.8407 | -100 | -100 | -100 | -52.3435 | -100 | -100 |
| **C5** | -35.4991 | -72.8879 | -5.12802 | -5.25052 | -0.0442 | -100 | -100 | -100 | -69.4189 | -100 | -53.4121 |

**Table S8.** Pdis-Msim EEAs for the 0h timepoint. Estimated community EEAs were calculated using the average EEAs of the constituent isolates. % proportional change of community EEAs was calculated as:
Community % change = (Measured community EEA - Estimated community EEA) / Estimated community EEA * 100

**Cleaved substrate concentration - Measured (µM min$^{-1}$)**

| **Isolate** | aG | bG | ACE | BUT | OLE | NAG | CB | XYL | PHOS | SULF | ARG |
|---|---|---|---|---|---|---|---|---|---|---|---|
| **AD-5** | 0.00622 | 0.05507 | 5.53802 | 3.13921 | 0.02237 | 0 | 0.01953 | 0 | 0.40969 | 0 | 1.53534 |
| **AD-12** | 0.01347 | 0.51613 | 5.56726 | 5.03677 | 0.00891 | 0.01293 | 0.06905 | 0.02725 | 0.15663 | 0.0017 | 0.03362 |
| **AD-55** | 0.15177 | 0.04177 | 0.56667 | 1.82224 | 0.02297 | 0 | 0.00482 | 0.0069 | 0.00731 | 0.00152 | 0.00071 |
| **AD-62** | 1.49079 | 0.61139 | 2.51396 | 1.27473 | 0.03096 | 0.01711 | 0.12876 | 0.00962 | 0.01263 | 0.00658 | 0.0115 |
| **AD-160** | 0.0357 | 0.09004 | 4.12216 | 3.17695 | 0.01757 | 0.00211 | 0.00373 | 0.00356 | 0.11499 | 0 | 0.20686 |
| **Community** | | | | | | | | | | | |
| **C1** | 0.38065 | 0.37561 | 3.77294 | 2.243 | 0.01502 | 0.00858 | 0.04084 | 0.00861 | 0.11242 | 0.003 | 0.21743 |
| **C2** | 0.05234 | 0.194 | 4.60783 | 3.17487 | 0.01483 | 0.00818 | 0.02915 | 0.00985 | 0.15785 | 0.00391 | 0.24672 |
| **C3** | 0.37484 | 0.31176 | 4.70327 | 2.96786 | 0.02709 | 0.00815 | 0.03918 | 0.00744 | 0.14539 | 0.00155 | 0.26752 |
| **C4** | 0.42366 | 0.29562 | 2.64871 | 1.93108 | 0.03062 | 0.0049 | 0.0445 | 0.00309 | 0.11028 | 0 | 0.23084 |
| **C5** | 0.4078 | 0.42711 | 3.55163 | 2.52719 | 0.02933 | 0.01354 | 0.05975 | 0.0113 | 0.06181 | 0.00468 | 0.07637 |

**Cleaved substrate concentration - Estimated community EEAs (µM min$^{-1}$)**

| **Community** | aG | bG | ACE | BUT | OLE | NAG | CB | XYL | PHOS | SULF | ARG |
|---|---|---|---|---|---|---|---|---|---|---|---|
| **C1e** | 0.41556 | 0.30609 | 3.54648 | 2.81823 | 0.0213 | 0.00751 | 0.05554 | 0.01094 | 0.14657 | 0.00245 | 0.39529 |
| **C2e** | 0.05179 | 0.17575 | 3.94852 | 3.29379 | 0.01795 | 0.00376 | 0.02428 | 0.00943 | 0.17216 | 0.0008 | 0.44413 |
| **C3e** | 0.38655 | 0.31816 | 4.43535 | 3.15691 | 0.01995 | 0.00804 | 0.05527 | 0.01011 | 0.17349 | 0.00207 | 0.44683 |
| **C4e** | 0.42112 | 0.19957 | 3.1852 | 2.35328 | 0.02347 | 0.00481 | 0.03921 | 0.00502 | 0.13616 | 0.00202 | 0.4386 |
| **C5e** | 0.42293 | 0.31483 | 3.19251 | 2.82767 | 0.0201 | 0.00804 | 0.05159 | 0.01183 | 0.07289 | 0.00245 | 0.06317 |

**% proportional change in community EEAs**

| **Community** | aG | bG | ACE | BUT | OLE | NAG | CB | XYL | PHOS | SULF | ARG |
|---|---|---|---|---|---|---|---|---|---|---|---|
| **C1** | -8.40127 | 22.7121 | 6.38562 | -20.4112 | -29.493 | 14.189 | -26.4692 | -21.2913 | -23.2988 | 22.3928 | -44.9957 |
| **C2** | 1.06492 | 10.3824 | 16.6975 | -3.61043 | -17.3984 | 117.611 | 20.0263 | 4.45125 | -8.30799 | 385.252 | -44.4483 |
| **C3** | -3.02929 | -2.00991 | 6.04071 | -5.9886 | 35.7494 | 1.35075 | -29.1011 | -26.3528 | -16.1949 | -25.1454 | -40.129 |
| **C4** | 0.60379 | 48.1317 | -16.8433 | -17.941 | 30.4704 | 1.94974 | 13.4845 | -38.4456 | -19.0053 | -100 | -47.3682 |
| **C5** | -3.5791 | 35.6633 | 11.2488 | -10.6263 | 45.9178 | 68.4716 | 15.8181 | -4.49963 | -15.2035 | 91.0845 | 20.8967 |

**Table S9.** Pdis-Mdis EEAs for the 0h timepoint. Estimated community EEAs were calculated using the average EEAs of the constituent isolates. % proportional change of community EEAs was calculated as:
Community % change = (Measured community EEA - Estimated community EEA) / Estimated community EEA * 100

**Cleaved substrate concentration - Measured (µM $min^{-1}$)**

| Isolate | aG | bG | ACE | BUT | OLE | NAG | CB | XYL | PHOS | SULF | ARG |
|---|---|---|---|---|---|---|---|---|---|---|---|
| **AD-4** | 0 | 0.12787 | 0.69553 | 0.62873 | 0.03225 | 0 | 0 | 0.00115 | 0.04529 | 0 | 0 |
| **AD-8** | 0.84516 | 0.07627 | 4.387 | 0.27457 | 0.09395 | 0.09051 | 0.00856 | 2.98979 | 0 | 0 | 0.03693 |
| **AD-13** | 0.03672 | 0.2862 | 2.4059 | 0.35824 | 0.07719 | 0 | 0.01414 | 0.06791 | 0 | 0 | 0 |
| **AD-31** | 5.01033 | 0.17859 | 4.65412 | 4.98252 | 0.15798 | 0.44478 | 0.0645 | 0 | 3.44959 | 0 | 0.04123 |
| **AD-34** | 0 | 0 | 1.68785 | 1.05084 | 0.99597 | 0 | 0 | 0 | 0 | 0 | 0 |
| **Community** | | | | | | | | | | | |
| **C1** | 1.2216 | 0.22425 | 2.39319 | 1.17349 | 0.10172 | 0.11347 | 0.04098 | 1.5313 | 0.82599 | 0 | 0.01831 |
| **C2** | 0.19376 | 0.07472 | 2.11983 | 0.88753 | 0.18724 | 0.01493 | 0.00518 | 0.07313 | 0.00559 | 0 | 0.00749 |
| **C3** | 1.43974 | 0.12194 | 3.34785 | 1.98198 | 0.19451 | 0.16 | 0.0194 | 0.12939 | 1.01199 | 0 | 0.02983 |
| **C4** | 1.04809 | 0.06432 | 2.08819 | 1.50034 | 0.22997 | 0.0994 | 0.01248 | 0.01102 | 0.86768 | 0 | 0.01194 |
| **C5** | 1.17295 | 0.08643 | 2.77969 | 1.48071 | 0.26194 | 0.11321 | 0.01033 | 0.06798 | 0.77394 | 0.00128 | 0.01874 |

**Cleaved substrate concentration - Estimated community EEAs (µM $min^{-1}$)**

| Community | aG | bG | ACE | BUT | OLE | NAG | CB | XYL | PHOS | SULF | ARG |
|---|---|---|---|---|---|---|---|---|---|---|---|
| **C1e** | 1.47305 | 0.16723 | 3.03564 | 1.56102 | 0.09034 | 0.13382 | 0.0218 | 0.76471 | 0.87372 | 0 | 0.01954 |
| **C2e** | 0.22047 | 0.12259 | 2.29407 | 0.5781 | 0.29984 | 0.02263 | 0.00568 | 0.76471 | 0.01132 | 0 | 0.00923 |
| **C3e** | 1.46387 | 0.09568 | 2.85612 | 1.73417 | 0.32004 | 0.13382 | 0.01827 | 0.74774 | 0.87372 | 0 | 0.01954 |
| **C4e** | 1.26176 | 0.14817 | 2.36085 | 1.75508 | 0.31585 | 0.1112 | 0.01966 | 0.01726 | 0.87372 | 0 | 0.01031 |
| **C5e** | 1.47305 | 0.13527 | 3.28372 | 1.66654 | 0.33127 | 0.13382 | 0.0218 | 0.76443 | 0.8624 | 0 | 0.01954 |

**% proportional change in community EEAs**

| Community | aG | bG | ACE | BUT | OLE | NAG | CB | XYL | PHOS | SULF | ARG |
|---|---|---|---|---|---|---|---|---|---|---|---|
| **C1** | -17.0699 | 34.0932 | -21.1636 | -24.8255 | 12.5951 | -15.2072 | 87.9793 | 100.245 | -5.46264 | - | -6.2667 |
| **C2** | -12.1141 | -39.0449 | -7.59527 | 53.527 | -37.5543 | -34.034 | -8.66842 | -90.4372 | -50.628 | - | -18.8399 |
| **C3** | -1.6483 | 27.4402 | 17.2166 | 14.2902 | -39.2234 | 19.557 | 6.21565 | -82.6962 | 15.826 | - | 52.6456 |
| **C4** | -16.9345 | -56.5865 | -11.5492 | -14.5147 | -27.1897 | -10.6075 | -36.5492 | -36.1435 | -0.69083 | - | 15.8238 |
| **C5** | -20.3727 | -36.1052 | -15.3492 | -11.1509 | -20.9299 | -15.4019 | -52.638 | -91.1066 | -10.2566 | - | -4.09579 |

**Table S10. Linear regression models for the effects of phylogenetic and metabolic diversity on the EEA proportional changes of bacterial communities.** Models include average EEA and EEAs for individual substrates. Significant effects are shown with their p values in **bold font.**

| | **Phylogenetic diversity** | | | | | **Metabolic diversity** | | | | | |
|---|---|---|---|---|---|---|---|---|---|---|---|
| ***Substrate*** | ***Estimate*** | ***± S.E.*** | ***t*** | ***p*** | ***df*** | ***Estimate*** | ***± S.E.*** | ***t*** | ***p*** | ***df*** | $R^2$ / $R^2$ ***adjusted*** |
| **Average** | 0.643 | 0.193 | 3.337 | **0.004** | 20 | -0.897 | 0.191 | -4.689 | **<0.001** | 20 | 0.701 / 0.645 |
| **aG** | 0.211 | 0.088 | 2.398 | **0.029** | 20 | -0.336 | 0.087 | -3.849 | **0.001** | 20 | 0.584 / 0.506 |
| **bG** | 0.368 | 0.251 | 1.463 | 0.163 | 20 | -0.760 | 0.250 | -3.046 | **0.008** | 20 | 0.430 / 0.323 |
| **ACE** | 0.038 | 0.048 | 0.802 | 0.435 | 20 | -0.007 | 0.047 | -0.153 | 0.880 | 20 | 0.162 / 0.005 |
| **BUT** | -0.095 | 0.063 | -1.509 | 0.151 | 20 | -0.053 | 0.063 | -0.839 | 0.414 | 20 | 0.289 / 0.156 |
| **OLE** | 0.020 | 0.089 | 0.229 | 0.821 | 20 | -0.227 | 0.088 | -2.576 | **0.020** | 20 | 0.340 / 0.216 |
| **CB** | 1.029 | 0.444 | 2.317 | **0.034** | 20 | -1.604 | 0.441 | -3.636 | **0.002** | 20 | 0.596 / 0.520 |
| **XYL** | 1.133 | 0.457 | 2.479 | **0.025** | 20 | -2.019 | 0.454 | -4.446 | **<0.001** | 20 | 0.620 / 0.549 |
| **NAG** | 1.470 | 0.455 | 3.231 | **0.005** | 20 | -1.828 | 0.452 | -4.046 | **0.001** | 20 | 0.646 / 0.580 |
| **PHOS** | 0.108 | 0.090 | 1.196 | 0.249 | 20 | -0.213 | 0.089 | -2.383 | **0.030** | 20 | 0.465 / 0.364 |
| **SULF** | -0.286 | 1.585 | -0.180 | 0.860 | 15 | -3.238 | 1.813 | -1.786 | 0.102 | 15 | 0.450 / 0.300 |
| **ARG** | 1.524 | 0.367 | 4.156 | **0.001** | 20 | -1.419 | 0.364 | -3.895 | **0.001** | 20 | 0.724 / 0.672 |
